# Single-cell Spatial Proteomic Imaging for Human Neuropathology

**DOI:** 10.1101/2022.03.02.482730

**Authors:** Kausalia Vijayaragavan, Bryan J Cannon, Dmitry Tebaykin, Marc Bossé, Alex Baranski, JP Oliveria, Dunja Mrdjen, M. Ryan Corces, Erin F McCaffrey, Noah F Greenwald, Yari Sigal, Zumana Khair, Trevor Bruce, Anusha Rajaraman, Syed A Bukhari, Kathleen S. Montine, R. Michael Angelo, Thomas J. Montine, Sean C. Bendall

## Abstract

Neurodegenerative disorders are characterized by phenotypic changes and hallmark proteopathies. Quantifying these in archival human brain tissues remains indispensable for validating animal models and understanding disease mechanisms. We present a framework for nanometer-scale, spatial proteomics with multiplex ion beam imaging (MIBI) for capturing neuropathological features. MIBI facilitated simultaneous, quantitative imaging of 36 proteins on archival human hippocampus from individuals spanning cognitively normal to dementia. Customized analysis strategies identified cell types and proteopathies in the hippocampus across stages of Alzheimer’s disease (AD) neuropathologic change. We show microglia-pathologic tau interactions in hippocampal CA1 subfield, in AD dementia. Data driven, sample independent creation of spatial proteomic regions identified persistent neurons in pathologic tau neighborhoods expressing mitochondrial protein MFN2, regardless of cognitive status, suggesting a survival advantage. Our study revealed unique insights from multiplexed imaging and data-driven approaches for neuropathologic analysis and serves as a baseline for mechanistic and interventional understanding in human neurodegeneration.

## INTRODUCTION

Neurodegenerative diseases are a heterogeneous group of disorders characterized by progressive dysfunction and loss of neurons, usually alongside aberrant protein accumulation. These proteopathies are accompanied by microglia (Keren-Shaul et al., 2017; Mrdjen et al., 2018) and astrocyte (Acioglu et al., 2021) stress responses and neurovascular dysfunction, and occur in characteristic brain regions that underlie clinical symptoms of disease. Alzheimer’s Disease (AD), the most common neurodegenerative disease, portrays these features. In AD, aggregation of amyloid-β (Aβ) peptides in parenchymal space as plaques and hyperphosphorylated, paired-helical filament (PHF) tau as intraneuronal neurofibrillary tangles (NFTs) and neuropil threads (NTs) form region-specific niches of neurodegeneration. Pathologic aggregates of PHF tau accumulate first in the entorhinal cortex and hippocampus; Aβ plaques deposit first across neocortical regions (Braak and Braak, 1991; Hyman et al., 2012).

Our understanding of the underlying phenotypic, regional, and cellular heterogeneity in neurodegenerative niches of these vulnerable brain regions remains limited(Mrdjen et al., 2019). Quantitative, multiplexed, subcellular imaging of cells and biochemical components may connect these various domains. Capturing discrete cellular and proteopathy features could be used to calculate local and regional tissue neighborhoods and their relationship to neurodegenerative disease progression. However, efforts to acquire highly multiplexed fluorescence-based images have been confounded by the variable auto-fluorescent matrix common to aged human brain formalin fixed paraffin embedded (FFPE) samples (Gilissen and Staneva-Dobrovski, 2013; Guardo, 2015). Furthermore, the ramified morphology of glia and neurons present significant analytical challenges to conventional methods for image segmentation and interpretation (Keren et al., 2019). Partitioning disjointed, unusually shaped or sized cellular features, acellular protein aggregates, and unclear tissue boundaries requires a specific set of analytical tools (Dora et al., 2017).

Here, we present a new generation of single-cell, spatial proteomic imaging, and analytical tools to capture the underlying cellular and phenotypic diversity in intricate neural tissues. In place of fluorophores and lasers, we employed multiplex ion beam imaging by time of flight (MIBI-TOF) mass spectrometry (MS) that has been used previously to study various archival human tissue types from tumors to placenta and granulomas in infectious disease (Greenbaum et al., 2021; Keren et al., 2018; Liu et al., 2021; McCaffrey et al., 2022; Risom et al., 2022). Based on secondary ion mass spectrometry (SIMS), MIBI-TOF images antigens targeted by antibodies labelled with elemental isotopic mass reporters that are spatially quantified with nanometer resolution. This MS-based strategy bypasses the light-based imaging matrix effects and autofluorescence that dominate in adult human CNS. Through verification against traditional IHC, we validated antibodies for 39 brain-specific targets (Table S1) and demonstrate these could be simultaneously stained and imaged with MIBI-TOF without the need for cyclic or serial approaches (see for comprehensive review of (Hickey et al., 2021). With these quantitative, multiplexed, spatial proteomic profiles across different brain regions, we were able to employ data-driven approaches to organize brain regions independently of local anatomy and disease status.

Finally, as a proof-of-utility for this methodological resource, we used archival human hippocampus from individuals covering a spectrum of cognitive impairment and AD neuropathologic changes. Adapting tools for automated feature identification, we extracted and classified a wide range of brain cellular features and protein aggregates. These were then combined to create tailored spatial analysis approaches, identifying organization within hippocampal subfields (Keren et al., 2018; Stoltzfus et al., 2020). Traditional single-cell data clustering, spatial correlation, and dimensionality reduction were used to reveal salient features. *Top-down* analysis, (leveraging prior knowledge of neuroanatomy), as well as *bottom-up* (completely data driven) approaches established complex spatial phenotypes across 275,808 cells and anatomical objects. Top-down analysis demonstrated changes in cellular composition across pathological stages related to AD progression, such as active microglia phenotypes in hippocampal subfield CA1 as evidenced by relatively high levels of IBA1, CD45, APOE, and CD33. Bottom-up analysis identified novel cellular neighborhoods with unique neuronal populations. Most interestingly, a pathologic Tau region present in the hippocampus of all individuals, regardless of cognitive status, showed a subset of persistent, pathologic protein free, neurons expressing increased levels of the mitochondrial protein MFN2. These insights were uniquely revealed by our multiplexed imaging and data-driving approaches tailored here for neuropathology research. Taken together, we provide a resource for acquisition, processing, and interpretation of highly multiplexed imaging of archival human brain tissue that should be extensible to platforms beyond MIBI-TOF presented here.

## RESULTS

### A framework for quantitative multiplexed imaging and feature selection in human brain

Archival, formalin fixed paraffin embedded (FFPE) brain regions are predominantly used for human neurodegenerative disease research. Fluorescence imaging on adult brain FFPE is confounded by endogenous tissue autofluorescence that can overwhelm signals originating from antibody bound fluorophores (Figure S1A) (Gilissen and Staneva-Dobrovski, 2013; Guardo, 2015; Schnell et al., 1999). With this in mind, we created a framework for high resolution, quantitative, multiplexed imaging of archival brain regions (Figure 1A) using MIBI-TOF. Because antibodies are detected using elemental mass tags, MIBI-TOF images possess no equivalent endogenous signal from the biological matrix (i.e., no equivalent ‘autofluorescence’). Brain tissue sections were stained with a cocktail of primary antibodies (36-plex panel), where each antibody is labelled with a unique elemental mass tag (Figure 1A, Figure 1SB, Table S1). Stained brain sections were imaged using an ion beam, which liberates these mass reporters as secondary ions (Figure 1A). The spatial distribution of elemental reporters is converted into *N*-dimensional image where each channel of this image corresponds to one of the primary antibodies (Figure 1A). Quantitative pixel-level information was extracted to produce global expression summaries and fed into downstream pipelines for data-driven analysis of cell phenotypes and disease objects classification *(see* Methods: *Global expression pattern*) (Keren et al., 2018). Ultimately, these data were used to construct tabular summaries of molecular features describing overall tissue architecture, as well as the spatial distribution of cells and proteopathies (Figure 1A).

**Figure 1.**
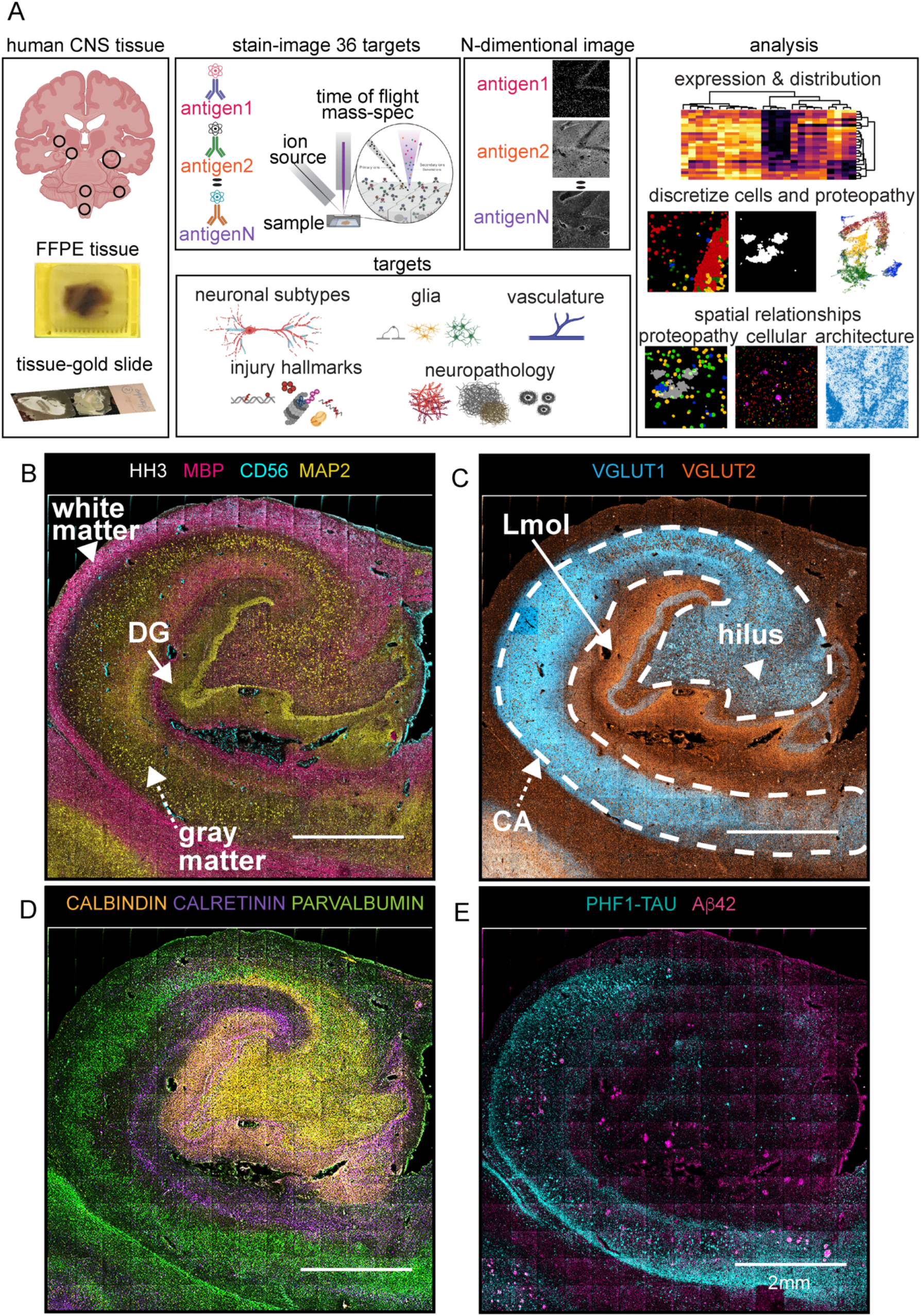
Workflow and Features Extracted from MIBI-TOF Spectral Images. **(A)** MIBI-TOF experimental workflow begins with the staining of FFPE brain regions (e.g., hippocampus, substantia nigra, striatum, locus coeruleus, medulla oblongata, cerebellum) on conductive gold sides, and imaging with a panel of 36 antibodies, each labelled with a unique elemental mass tag. Stained tissue sections were rasterized and resulted in a 36-dimensional image revealing protein expression at unique spatial coordinates in a selected field of view (FOV). MIBI-TOF generates data to quantify and visualize tissue architecture, which allows analysis of global protein expression and distribution, discretization into biological units for exploring marker relationships within the biological units, and investigation of distance relationships and neighborhoods among these biological units. **(B-E)** Montage of 196 tiled FOVs MIBI-TOF image from a post-mortem FFPE archival human hippocampus from an individual with AD dementia (ADD). Each tile is a 500μm^2^ spectral image acquired at low-resolution scan resolving ∼1μm^2^/ pixel. Representative 11 out or 36 spectral images were pseudo-colored and overlaid to show the spatial distribution and expression of structural and molecular markers. Abbreviations: DG, Dentate Gyrus; CA, Conus Ammonis; Lmol, molecular layer.

To capture salient neuropathologic changes, the antibody staining panel targeted proteins for delineating major neuronal cell lineages, proteopathies, and vascular structures (Figure S1B, Table S1). Antibody specificities were validated against multiple brain regions by standard IHC (Figure S1C), benchmarking optimal titers of chromogenic IHC against MIBI-TOF. We used the resulting 36-plex panel to reveal the cytoarchitecture of an entire coronal section of human hippocampus (Figure 1B-E). Hippocampus consists of anatomically distinct regions with structural boundaries that are evident from pseudo-colored overlays. Figure 1B-E illustrates a 31mm^2^ hippocampal region from a person with AD dementia (ADD). The granule cell layer of dentate gyrus (DG) is highlighted by MAP2 and HH3 (Figure 1B, DG *white arro*w), while the boundary between white and gray matter can be delineated based on expression of MBP and CD56 (Figure 1B). In line with previous work, glutamatergic terminals within the hilus, Cornu Ammonis (CA), and molecular layers (Lmol) are marked by the presynaptic vesicular proteins VGLUT1 and VGLUT2 (Figure 1B) (Herzog et al., 2006; Liguz-Lecznar and Skangiel-Kramska, 2007; Vigneault et al., 2015). Calbindin (CB), calretinin (CR), and parvalbumin (PV) appear as contiguous parcels in the hippocampus (Figure 1D). As seen in rat hippocampus, our human data showed that CB borders the DG and hilus, CR structures the Lmol, and PV is within CA (Rivera et al., 2014). As expected, NFTs and NTs are highlighted by PHF1-TAU immunoreactivity in the CA subfield, while Aβ plaques depicted by Aβ42-immunoreactivity predominantly in hilus and Lmol of the ADD hippocampus (Figure 1E). Spectral images of all simultaneously acquired markers are illustrated in Figure S4I. Taken together, this comprehensive survey of brain phenotypes highlights the capability of our approach to visualize multiplexed signatures that reveal salient structural hallmarks and proteopathies in human hippocampus.

### Multiplex imaging protein signatures organize anatomical structures

Given the multiplexed signatures that demarcated known subfields within the hippocampus, we sought to determine how they might quantitatively organize brain cytoarchitecture across a broader set of regions (Figure 2). We used our 36-plex panel to analyze a tissue microarray (TMA) containing six different brain regions from cognitively normal individuals (Figure 2B). Many of these markers exhibited noticeable gradients that were concordant with canonical multicellular structures. For example, calcium-binding proteins (PV, CB and CR) show laminar distributions and densities that are consistent with previously reported protein and gene expression in mouse and human, within and across these brain regions (Figure 2B, Figure 2C) (Bjerke et al., 2021; Hawrylycz et al., 2012; Hof et al., 1999). With the exception of the medulla oblongata (MO), unsupervised clustering of extracted cytoarchitectural features alone organized images separated based on their respective brain regions (Figure 2B). We also used this approach to delineate features from individuals with cognitive impairment (Figure S2B). Taken together, much like multiplexed analysis of the immune system and other tissues (Glass et al., 2019; Hartmann and Bendall, 2020; Hartmann et al., 2021; Keren et al., 2018; Liu et al., 2021; McCaffrey et al., 2022; Moore et al., 2021), combinatorial protein expression patterns provided a snapshot of functional organization or proteopathy within brain regions. This low-level analysis of multiplexed proteomic data could serve as a guide for ‘fingerprinting’ human brain and be used to model progression in neurodegeneration.

**Figure 2.**
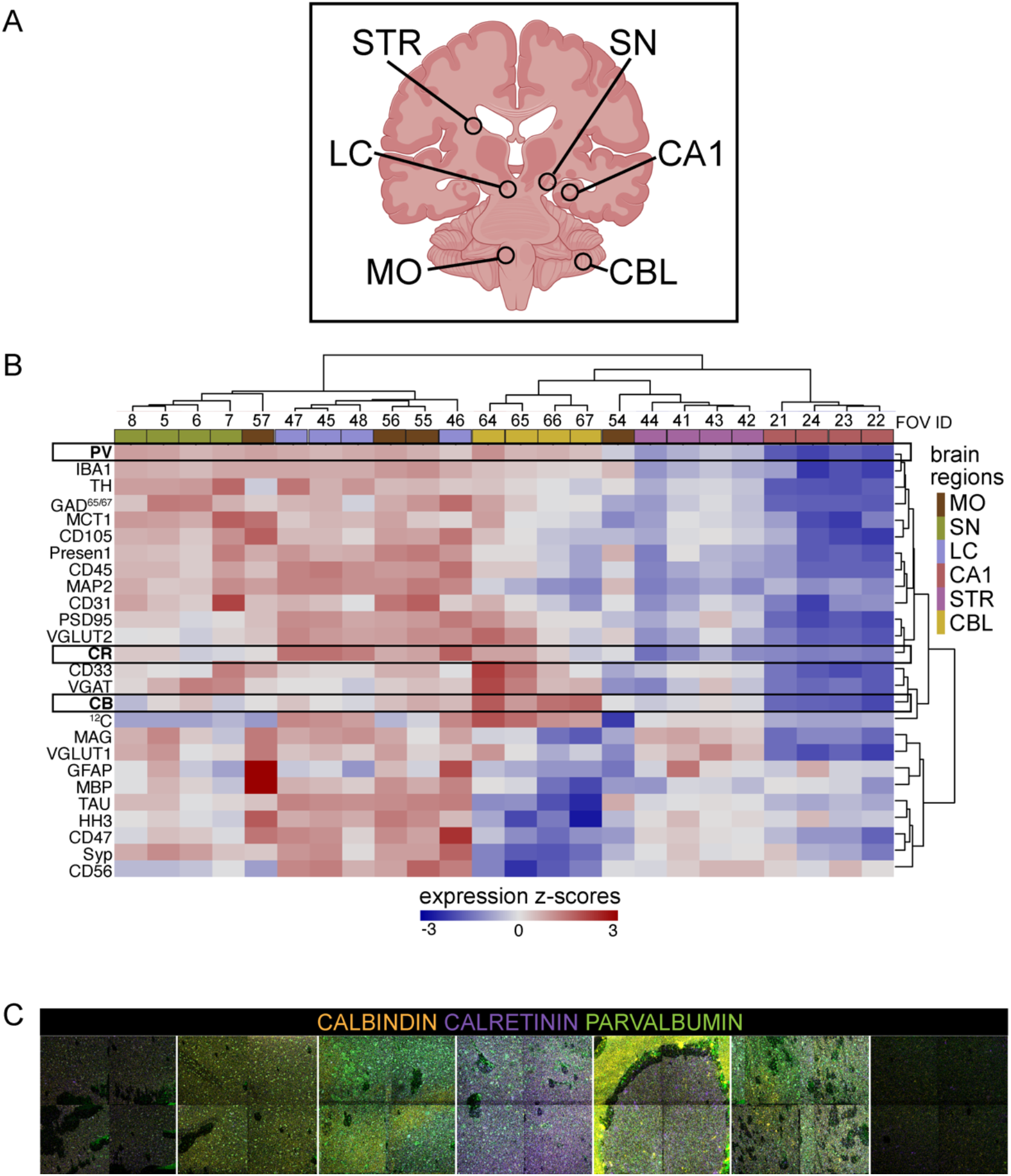
Global phenotypic expression organizes CNS sub-regions in a data driven manner. **(A)** Schematic of brain regions used in construction of TMA; circle highlights 3mm cores isolated from FFPE tissues. **(B)** Heatmaps showing the z-score distribution of pixel expression of proteins per FOV. Columns and rows are hierarchically clustered (Euclidean distance). Variance between and among FOVs are stratified in an unsupervised manner into similar anatomical regions. **(C)** Spectral image pseudo-colored to show distribution of calcium-binding protein PV, CB and CR. Abbreviations: STR, Striatum; LC, Locus Coeruleus; MO, Medulla Oblongata; SN, Substantia Nigra; CA1, Cornu Ammonis 1; CBL, Cerebellum; CB; CALBINDIN; CR, CALRETININ; PV, PARVALBUMIN.

### Discretized cellular and pathological features identify lineage and disease pathology-specific subclusters

We next discretized cells and proteopathies as biological units: neurons, astrocytes, microglia, vasculature, NFTs-NTs, and Aβ plaques. Segmentation of planar brain tissue sections has inherent problems due to shape, texture, the disjointed features from neuronal and glial processes in the tissue section, but distant from their cell bodies, and from nonconforming shapes from protein aggregates. As shown in Figure 3A (*dashed lines*), we observed these various facets of planar imaging by both standard IHC and MIBI-TOF. To capture these attributes, two segmentation methods were used to partition neuronal perikaryons, microglia, astrocytes, and their processes, endothelial and their vascular-boundaries, Aβ plaques, and NFTs-NTs (Figure 3B). An adapted version of DeepCell was used for nuclear segmentation (Figure 3B) (Greenwald et al., 2021; Moen et al., 2019; Valen et al., 2016). For features not associated with nuclei, a pixel intensity thresholding-based method was used to partition cell body microglial and astrocyte projections, larger vascular structures, amyloid plaques, and NFTs-NTs (Figure 3B, see Methods: *Object segmentation*).

**Figure 3.**
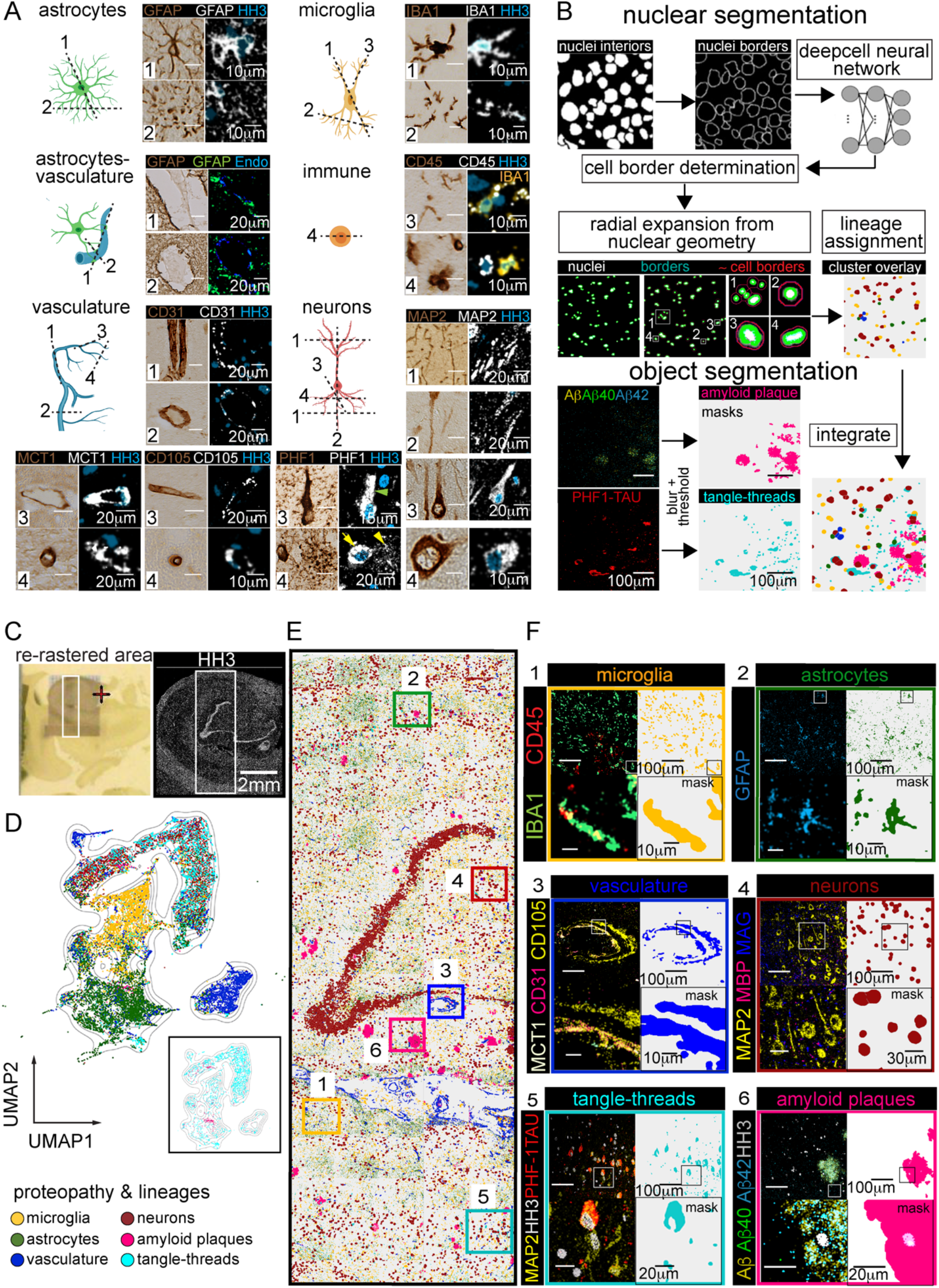
Spatial Organization and Molecular Identities of Cells and Proteopathy in ADD hippocampus is Revealed with Single-Cell and Object Segmentation. **(A)** Cartoons show examples of cells (astrocytes, microglia, vessels, neurons, and immune cells) and possible planar views in CNS tissue (dashed lines numbered 1 through 4). The representative photomicrograph (IHC-DAB) and MIBI-TOF (pseudo-colored spectral images) visually exemplify the corresponding focal planes (numbered 1 through 4) for the different cells, which are identified by their pan-markers (GFAP: astrocytes, IBA1-CD45: microglia, CD31-CD105-MCT1: vasculature, MAP2: neurons, CD45: immune cells). **(B)** Conceptual overview of nuclear and object segmentation approaches. With cellular segmentation, nuclear associated features (nuclei interiors, nuclei borders) are obtained using a pixel-based convolution neural network (DeepCell neural network) and custom pixel expansion around the nuclei, to generate cell segmentation masks for cell lineage assignments. Object segmentation was used to capture microglia and astrocytic processes, vascular boundaries, Aβ plaques, and PHF1-TAU NFTs and NTs. Masks for both discretized biological units are integrated to reveal their relationships *in silico*. **(C)** Photomicrograph (on the conductive gold slide, left panel) and corresponding MIBI-TOF spectral image (HH3 for nuclei, right panel) of human ADD hippocampus. White box highlights the re-rastered region that was used for analysis. **(D)** UMAP visualization of all segmented objects across all hippocampal FOVs, colored by lineage and proteopathy identities. Inset shows only amyloid plaques, and tangles and threads overlaid on the UMAP. **(E)** Cell Phenotype Maps (CPM) of nuclear and object segmented masks labelled by their phenotype. **(F)** Composite CPM of all segmented masks with six inserts for a zoomed in view of cell identity overlaid onto segmentation masks for the hippocampal region. Regions 1-6 represent zoomed in images representative areas of CPM’s in E.

The same 36-plex stain of ADD hippocampus from Figure 1B was chosen for the discretization analysis to capture a greater breadth of morphological and pathological landscapes. With nuclear segmentation, we identified 15,270 nuclei-associated cells across 85 connected imaging FOVs which were classified using manual gating (Table S4, *see* Methods: *Manual gating ADD hippocampus*). With the pixel-based approach, we identified 19,332 features not associated with nuclei that included microglia, astrocytes, endothelial cells, Aβ plaques, and NFTs-NTs (Table S4). Neuron masks identified by nuclear segmentation were integrated with masks of the object segmented data (microglia, astrocytes, vasculature, plaques, and NFTs-NTs) to obtain a comprehensive repertoire of neuronal and non-neuronal cell types as well as disease features. We then generated a UMAP (McInnes et al., 2018) plot organized by lineage and proteopathy markers to analyze the relationship between these features (*see* Methods:*1024 x1024 ADD Images*). Extracted cellular and proteopathy features mapped to four cell-type clusters (neurons, microglia, astrocytes, and vasculature) and two pathologic component clusters (Aβ plaques and NFTs-NTs) (Figure 3D, inset).

NFT-NT masks (cyan) mostly interrelated with the neuron (dark red) cloud (Figure 3D, Figure S3K). This is consistent with known accrual of NFTs-NTs in intraneuronal compartments (Mrdjen et al., 2019; Uchihara, 2014). Aβ plaques shared the UMAP space mainly with astrocyte (dark green, Figure 3D, Figure S3L) and neuronal populations (Figure 3D, Figure S3M) as well as scattered with microglia (dark yellow) and vascular groups (Figure 3D). In accordance with previous reports, both astrocytes and neurons highly express Aβ, and GFAP+ reactive astrocytes accumulate Aβ in the process of clearance in AD (Calhoun et al., 1999; Wang et al., 2020; Zhou et al., 2017).

By mapping the masks of these gated objects back onto the coordinates of the original segmented images (Figure 3E), we created a cell phenotype map (CPM) that illustrated a more even distribution of hippocampal microglia relative to astrocytes (Figure 3E, Figure 3F1, Figure S3A). Astrocytes were enriched in hippocampal white matter and vascular regions (Figure 3E, Figure 3F2, Figure S3B) as they are integral parts of CNS white matter and blood-brain-barrier (BBB) architecture (Heithoff et al., 2020; Liedtke et al., 1996). Vascular CPM charts large and micro vessel boundaries (Figure 3E, Figure 3F3, Figure S3C). In the neuron CPM, masks are enriched in the granule cell layer of DG band, and are scattered throughout the image (Figure 3E, Figure 3F4, Figure S3D). These data highlight the utility of the combined segmentation approaches, enabling analyses analogous to those achieved in other tissue where spatial organization, expression, and molecular identities of single cells and proteopathy can now be determined and organized for human CNS.

### *Top-down* spatial organization of cellular and proteopathy composition distinguishes hippocampal subregions and disease status

We combined our discretization strategies with prior knowledge of human hippocampal anatomy (i.e., top-down) to identify neuropathologic changes across stages of AD. Coronal hippocampal sections from three individuals were imaged, focusing on the DG to CA4 through CA1 subregions, and expanding outward (Figure 4). Samples from cognitively normal (CN), cognitively impaired with no dementia (CIND), and ADD subjects were used to capture a wide spectrum of pathologic changes (Figure 4A). Single-plex spectral images of CN (Figure S4A-C), CIND (Figure S4D-F) and ADD (Figure S4G-I) hippocampus demonstrated larger, qualitative structures that can be captured by tiling numerous imaged fields together. For instance, VGLUT1 and VGLUT2 positivity demarking the CA and DG borders, respectively.

**Figure 4.**
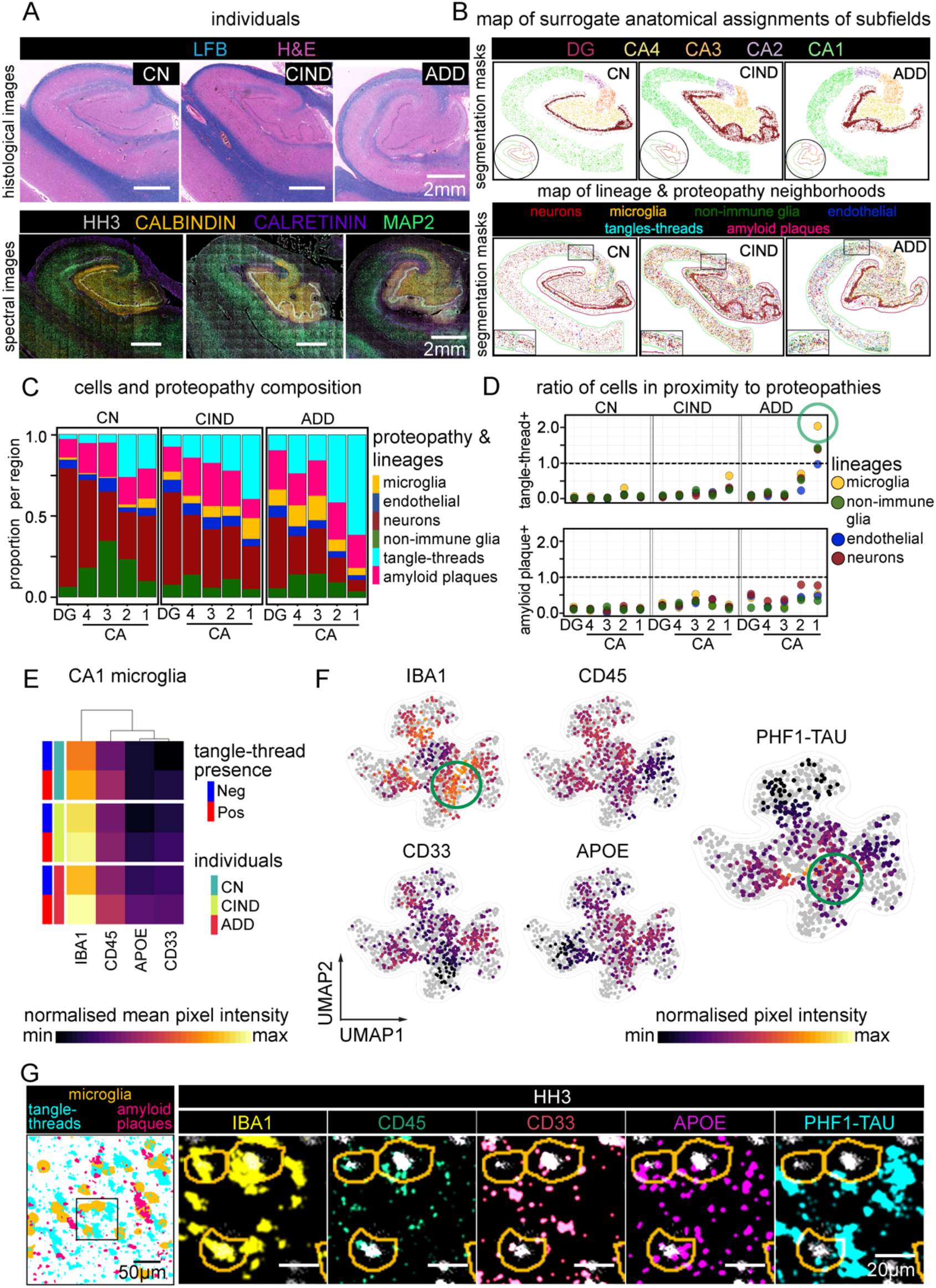
Cellular and proteopathy composition of hippocampal neuroanatomy. **(A)** Representation of the three imaged hippocampi to identify differences in AD pathological changes. Included are samples from people who were diagnosed as cognitively normal (CN), cognitively impaired, no dementia (CIND), or Alzheimer’s Disease dementia (ADD). LFB & H&E histochemical staining of a serial section of each hippocampal slice imaged (Top). Pseudo-color overlay of CALRETININ (CR), CALBINDIN (CB), MAP2, and Histone H3 (HH3) acquired in each sample representing high-level neuronal morphology (Bottom). **(B)** Mask overlays of the cells and proteopathies collected by the segmentation methods described in Figure 2, broken down by hippocampal neuroanatomy (Top) and cell or proteopathy subtype (Bottom). **(C)** Relative composition of each cell and proteopathy subtype within each anatomical subregion of each individual. **(D)** Ratio of proteopathy-associated cells to proteopathy-free cells in each anatomical region of each individual. Ratio of 1 indicates an equal number of proteopathy-associated cells to proteopathy-free cells, where PHF1-TAU tangle-thread cell associations are shown in the top-panel and Aβ plaque cell associations in the bottom-panel. Callout indicates ADD, CA1 microglia with a high number of tangle-thread-associated microglia (green circle). **(E)** Normalized mean expression of microglia phenotyping channels in CA1 region of each individual, broken down by microglia with tangle-thread association or those without. **(F)** UMAP projections of CA1 microglia phenotyping channels in ADD. Gray represents CA1 microglia from CN, CIND samples (S4K-L for CN, CIND expression maps). Projection was calculated using phenotypic markers. CIND and ADD are subsampled so that all conditions are represented by 447 microglia, the total number of CA1 microglia in the CN condition. **(G)** Image representation of tau-tangle associated phenotypes described in E and F found in ADD CA1 microglia. Masks of microglia (yellow), tau tangles (cyan), and amyloid plaques (pink) (Left). Inset of microglia expressing tau-tangle associated phenotypic markers (Right).

Given the known differences and progression of disease within AD hippocampal subregions (Braak and Braak, 1991), we first segregated all cells and disease objects into DG, CA4, CA3, CA2, and CA1 associations (Figure 4B). We determined these boundaries for the subregions based on prior knowledge of morphological characteristics (Lorente-de-Nó, 1934; RamonYCajal, 1902) and expression of neuronal makers including CALBINDIN, CALRETININ, MAP2, CD56, SYNAPTOPHYSIN (SYP), VGLUT1, and VGLUT2 (Figure 4A, Figure S4) that have previously been shown to highlight boundaries (Eriksson et al., 1998; Herzog et al., 2006; Rivera et al., 2014). Spectral images of CALBINDIN, CALRETININ, and MAP2, specifically showed anatomical delineation between the inner regions of DG and CA regions (CALBINDIN) and distal regions (CALRETININ), joined by neuronal cytoarchitecture (MAP2) (Figure 4A, bottom panel). We then assigned cells and protein aggregates to their respective structures (Figure 4B). (Table S5A). To assign cell identities, we applied a combination of FlowSOM (Gassen et al., 2015) and manual meta-clustering strategies to parse out the objects into six distinct categories: neurons, microglia, endothelial cells, and non-immune glia which included both astrocytes and oligodendrocytes (Figure 4B, lower panel).

Comparing the proportion of each cell type within each distinct anatomical region, we observed a higher proportion of protein aggregates in the CA1 subfield relative to other regions within the same sample, regardless of cognitive status (Figure 4C). In addition, the total number of disease objects increased in the tissue prom cognitively impaired individuals (Figure S4J). We then considered whether proteopathies tended to lay in proximity to cell somas bound by our cellular segmentation. For each cell subtype, we counted the number of cells in direct proximity (i.e., overlapping pixels) to each proteopathy and divided this value by the number of cells with no proteopathy overlap (Figure 4D). Most cells did not exhibit direct proximity to Aβ plaque objects, particularly in the CN and CIND conditions. In the ADD individual, more cells lay in proximity to Aβ plaques, particularly for neurons in the CA1 and CA2 subfields. For PHF1-TAU labeled NFTs-NTs, a similar trend was found, though interestingly ADD CA1 showed increasing proximity for all cell types. In particular, microglia with Tau aggregates were found twice as much as microglia without Tau aggregates in the ADD CA1 subfield (Figure 4D, *green circle*).

Building upon this observation, we contrasted those microglia that were NFTs-NTs associated (positive) or not (negative) (Figure 4E). Microglia markers associated with reactivity, APOE, IBA1, CD33, and CD45, were consistently higher in tangle-positive relative to tangle-negative CA1 microglia, across all samples, with the highest expression in ADD (Figure 4E). Projecting an equal subset of microglia from all three samples onto an 2D-UMAP embedding, shows subsets of these reactive microglia associated with PHF1-TAU, particularly high IBA1(+) and APOE(+) in CA1 ADD cells (Figure 4F, *green circle*); and that while these cells exist across all individuals, they are most prominent in ADD (Figure 4F, S4K, L). This differential state of expression further reflects the possibility that immune cell reactivity may play a central role in AD severity, particularly an association with PHF1-TAU formation in CA1 subregion.

### *Bottom-up*, data driven neighborhood analysis identifies common regions of neuropathology across individuals and severity

Given the level of neuropathological organization identified using previously defined hippocampal subregions, we investigated what other levels of spatial order could be revealed with a more data-driven approach to our multiplexed images. To this end, we employed a *bottom-up* workflow to isolate common signatures of hippocampal spatial identities, independent of previously known neuroanatomy or cognitive status. Using CytoMAP (Stoltzfus et al., 2020), we grouped local neighborhoods of similar protein aggregates and cell types across the three samples. After neighborhoods were calculated, a minimum number of common regions into which they could be grouped into was determined (*see* Methods: *Object co-proximity analysis*). Identified cells and protein aggregates were then assigned to these regions across each sample. A voronoi expansion from cell and object bearing areas was then used to cluster the surrounding neuropil (i.e., cell projections and extracellular matrix) into one of the regional groups (Figure 5A). The final five regions determined across all samples were annotated as neuronal dominant (N), glial dominant (G), mixed disease (M), Aβ plaque dominant (AP), or PHF1-TAU NFTs-NTs dominant (TT) (Figure 5B). The three proteopathy associated regions: M, AP, and TT were increased in representation with increasing cognitive impairment, while the relatively protein aggregate-free neuronal (N) and glial (G) cell regions decreased with increasing cognitive impairment, making up less than a quarter of the total ADD tissue (Figure 5C). As has been described extensively, even the cognitively normal hippocampus contained AP and TT regions (Figure 5B, left purple and blue), consistent with the high prevalence of pre-clinical AD in older people. Moreover, due to the data-driven nature of their identification we saw similar compositions of cells and disease objects within a given region and across samples, regardless of cognitive diagnosis (Figure 5D).

**Figure 5.**
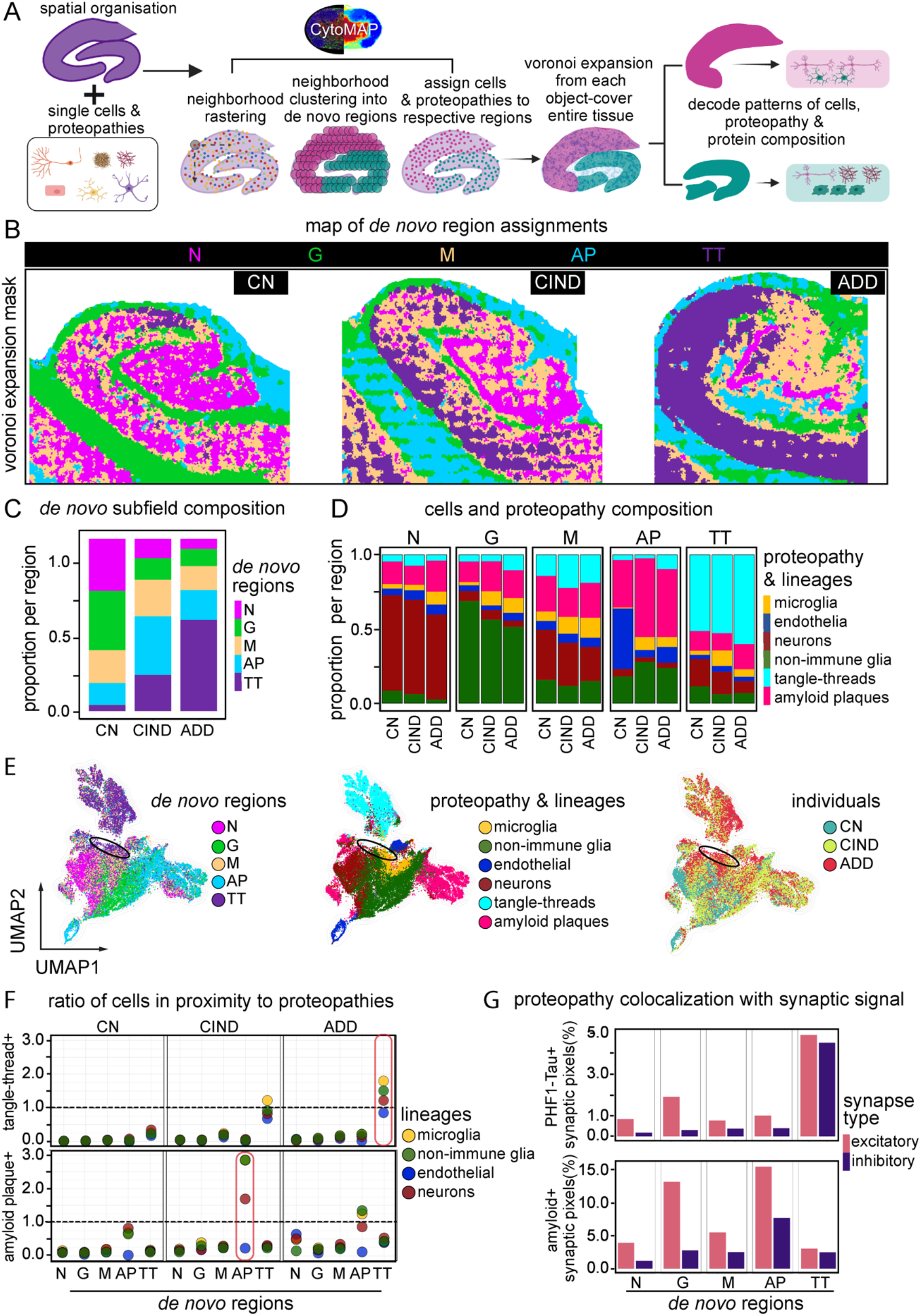
Common *de novo* regions of cell and proteopathy composition are identified across different individuals. **(A**) Computational workflow used to isolate common regions of pathology (*de novo* regions) across CN, CIND, and ADD hippocampi independent of patient diagnosis. **(B)** *De novo* regions displayed across the entire tissue of each sample, CN, CIND, and ADD. N=Neuronal dominant region, G=glial (astrocyte/oligodendrocyte) dominant region, M=Mixed proteopathy region, AP=Amyloid Plaque dominant region, TT=Tau Tangle-Thread dominant region. **(C)** Relative composition of each *de novo* region across each individual hippocampus. **(D)** Relative composition of each cell and proteopathy subtype within each *de novo* region of each individual. **(E)** UMAP projection of all cells and proteopathies, calculated using much of the full panel (list in methods). *De novo* regions (left), cell and proteopathy classification (middle) and individual (right). Black ellipse denotes area of clustered cell lineages associated with TT region. **(F)** Ratio of proteopathy-associated cells to proteopathy-free cells in each *de novo* region of each individual. Ratio of 1 indicates an equal number of proteopathy cells to proteopathy-free cells. Tau tangle cell associations (Top) and amyloid plaque cell associations (Bottom). Top callout indicates high levels of tau tangle overlap in TT region; bottom callout indicates high amyloid plaque overlap in AP region. **(G)** Synaptic proteopathy presence. For each de novo region, all positive synaptic pixels with positive proteopathy signal (Top: tau, Bottom: amyloid) relative to all positive synaptic pixels. Excitatory synaptic channels (SYP+, PSD95+, and either VGLUT1+ or VGLUT2+) and inhibitory synaptic channels (SYP+, VGAT+, GAD+) considered.

To integrate these regions and pathologic changes into a single model, we applied UMAP (McInnes et al., 2018) dimensionality reduction to all cells and protein aggregates, comparing how they overlapped in disease severity or data-driven regions (Figure 5E). While like cells or protein aggregates clustered tightly, there was some divergence with *de novo* region, suggesting unique combinations of protein expression with both anatomical location and spatially associated pathologic features. For example, a subset of neurons, non-immune glia, and microglia clustered with the TT region, separate from those in the N or G regions (Figure 5E, *black circle*). Given the influence of *de novo* identified regions on cell and object embedding (Figure 5E), we also quantified how cell-protein aggregate proximity itself compared between the top-down anatomical (Figure 4D) and data-driven regions (Figure 5F). Regardless of sample staging, cells within the N- or G-regions showed very little proximity to pathologic objects, as expected. In the AP-region, nonimmune glia and neurons showed high proximity to neurons and nonimmune glia (Figure 5F). The same trend held for PHF1-TAU NFTs-NTs in proximity to cells in the TT-region. As pathologic changes increased, microglia once again stood out with the highest ratio of cells with NFTs-NTs proximity. These physical associations increased as severity worsened whether the nuclear associated cell body or protein aggregates were used as the basis for the proximity calculation (Figure S5A). In summary, while NFTs-NTs development started in neuronal peripheries before accumulating in soma (Brunello et al., 2020; Tai et al., 2014; Uchihara, 2014) our data suggests soma involvement continues as disease progresses.

With the connection between proteopathy formation and synaptic density, we further investigated the mean pixel expression of these markers across all identified data-driven regions in the hippocampus and how coincidence changed. Focusing on synaptic density, we calculated synaptic positive pixels by either excitatory (SYP+, PSD95+, and either VGLUT1+ or VGLUT2+) or inhibitory (SYP+, VGAT+, GAD+) synaptic pixels. We then looked at the coincidence with PHF1-TAU and observed that while the highest percentage of PHF1-TAU formation in synapses occurred in ADD, all samples that contained the TT-region showed synaptic protein/PHF1-TAU localization (Figure 5G), similar to what we have recently reported in synaptosomes (Phongpreecha et al., 2021). Similar results were seen for Aβ plaque proximity to synaptic pixels in the AP region (Figure 5G). This approach, as summarized in Figure 5A, shows the ability to identify common pathological changes and neighborhoods across tissues representing a diversity of neuropathologic changes. Moreover, completely data-driven, bottom-up approaches, like the ones employed here, captured, and organized neuropathological features beyond the bounds of conventional anatomical subregions potentially increasing the sensitivity to detect and classify dysfunction.

### Clustering identifies mitochondrial MFN2 protein expression in persistent neurons associated with proteopathy-laden tissue regions

Leveraging the 14 proteins simultaneously measured to capture neuron identity and putative function *in situ*, we clustered these extracted cells using FlowSOM to identify 11 meta clusters. The expression distributions across these groups (Figure 6A) and as seen by UMAP projection (Figure 6B-C) and physical mapping (Figure S6A) showed that neurons fell broadly into two categories: 1) proteopathy associated (clusters 1, 2, 6, 8, 10), and 2) non-proteopathy associated (clusters 3, 4, 7, 9, 11). While this analysis highlighted the previous trend of increased protein expression in proteopathy-associated cells (i.e., synaptic markers, Figure 6A, S5B) it revealed some functional neuronal subsets based on mitochondrial proteins and ubiquitination, e.g., MITOFUSION 2 (MFN2) (Figure 6A, clusters 1, 2, 3, 4, 10) and POLYUBIQUITIN K48 (Figure 6A, clusters 1, 2, 8).

**Figure 6.**
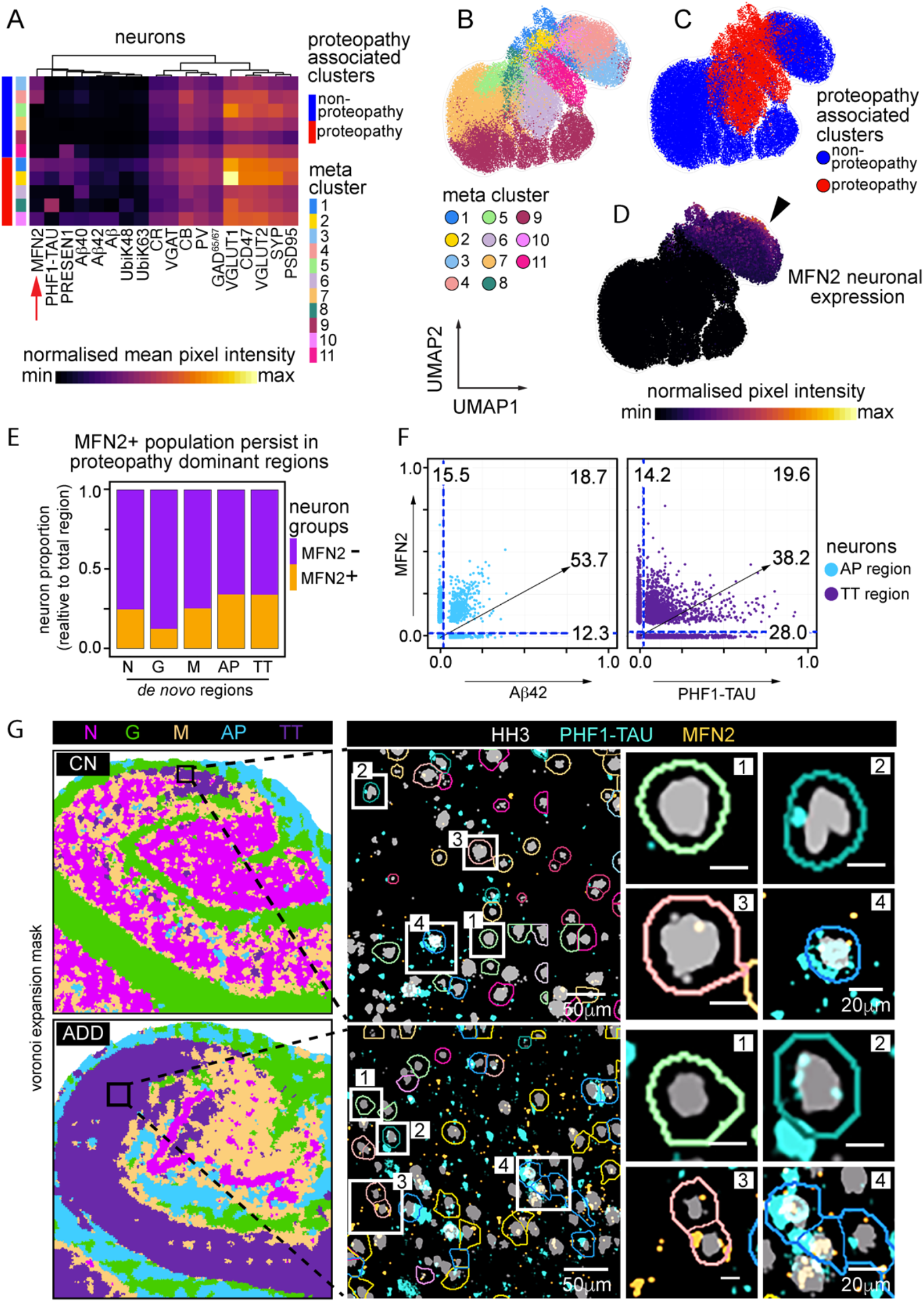
Neuronal sub-clustering reveals relationship between MFN2 and persistent neurons in AD. **(A)** Heatmap showing mean normalized expression of neuronal markers after FlowSOM clustering. 11 clusters are broadly broken down into proteopathy and non-proteopathy associated phenotypes. Red arrow denotes MFN2 cluster expression. **(B-D)** UMAP projection of neurons alone, calculated based on same markers used to perform FlowSOM clusters in **A. (B)** Overlay of neuronal subclusters. **(C)** Overlay of clusters associated with proteopathy or no-proteopathy (D) Overlay of normalized neuron expression of MFN2. **(E)** Proportion of neurons positive for MFN2 vs neurons negative for MFN2 for each *de novo* region. **(F)** Expression profile of individual neurons for MFN2 and Aβ42 in AP region (left) and MFN2 and PHF1-TAU in TT region (right). **(G)** Neurons in TT region of CA1, CN individual (top) and CA1, ADD individual (bottom). Black box represents areas of TT-associated tissue isolated for insets (left), zoomed in section of multiple neurons, colored by cluster ID’s in A (middle), single or small groups of neurons representing persistent, TT cells (right). [1] MFN2(-), PHF1-TAU(-) neuron – Cluster 5. [2] MFN2(-), PHF1-TAU(+) neurons – Cluster 8 [3] MFN2(+), PHF1-TAU(-) neurons – Cluster 4 [4] MFN2(+), PHF1-TAU(+) neurons – Cluster 1.

As with the broader analysis of all objects (Figure 5), neuron subclusters associated with protein aggregates grouped together in a UMAP of neurons alone (Figure 6C, left). As MFN2 was variably expressed across multiple non-proteopathy associated (3, 4) and proteopathy-associated neuronal clusters (1, 2, 10), we further investigated its expression and localization in the hippocampus. Interestingly, recent single nucleus analysis of transposon accessible chromatin (ATAC-seq) indicates that MFN2 is epigenetically restricted to neuron-specific lineages in human brain (Figure S6B) (Corces et al., 2020), re-enforcing its potentially unique role in neuronal function here. Neuronal MFN2 expression was positive primarily in non-proteopathy associated neurons, with a clear overlap in proteopathy associated neurons. (Figure 6D, *right, black arrow*). These MFN2 positive neurons were more prevalent in proteopathy-associated regions (MD, AP, TT) relative to the non-proteopathy associated regions (ND, GD) (Figure 6D), suggesting that MFN2 expression may be associated with neuronal survival in the presence of stressors from proteopathy.

To understand MFN2-related persistence in proteopathy-laden regions, we compared the single-cell relationship between proteopathy expression and MFN2. Filtering by *de novo* region shows high abundance of MFN2(+), PHF1-TAU(+) neurons in the TT region (∼20%) compared to the N region (∼1%) (Figure 6F, *right*, S6C). These PHF1-TAU(+), MFN2(+) neurons can be seen throughout the TT region of the hippocampus, even appearing adjacent to PHF1-TAU single positive, as well as MFN2 single positive expressing neurons, regardless of disease status (Figure 6F). Comparatively, the N region contained many more MFN2+ single-positive neurons. This persistence of MFN2+ neurons in areas of high tauopathy potentially reflects a neuronal protective response to injury as has been studied in animal and *in vitro* models of neuronal degradation (Motori et al., 2020). MFN2’s relationship with Aβ42 did not show this same L-shaped distribution, especially in the AP region (Figure 6F, *left*), suggesting that the presumed MFN2 response to injury may be specific to stressors that accompany pathologic Tau accumulation. Altogether, our study demonstrates both anatomical and data-driven approaches can be used to identify deep spatial summaries of neuropathologic changes while identifying and subclassifying cells and features that compose them.

## DISCUSSION

In this study, we present an analytical framework leveraging MIBI-TOF spatial proteomic imaging as a resource for deep human neuropathology. We describe a workflow for the 36-plex image datasets generated from archival FFPE human brain tissues. We performed a broad IHC analysis that validated MIBI-TOF spectral images in multiple brain regions with neuropathologic changes spanning different stages of AD both clinically and pathologically. Our imaging panel was designed to phenotype brain cellular, structural, and proteopathy features while displaying well-known brain topologies. Segmentation of single-cell and disease features were phenotypically profiled, and their spatial organization in relation to local and global neighborhoods were explored. Finally, to delineate the cellular and molecular configuration in different clinic-pathological states of human AD, we showed the effectiveness of complementary approaches: a ‘top down’ method, that relied on classically stratified hippocampus subregions (DG through CA4-CA1), and a bottom-up data driven method that defined proteopathy-free and proteopathy-burdened cellular neighborhoods within the human hippocampus.

Notably, our study represents one of the most systematic, high-resolution analyses of *spatial* cellular and proteopathy composition in human brain. Age- (98.3 ± 1.5 yrs.), sex- (female), and *APOE* genotype- (epsilon3/epsilon3) matched samples were compared across carefully annotated clinico-pathologic samples from three research participants. We primarily focused on changes in the hippocampus, due to its implication early in AD pathogenesis (Braak and Braak, 1991). We have configurations for the abundance of 34-plex proteins and their spatial distributions across 275,808 segmented cells, 25,130 Tau protein aggregates and 15,174 amyloid protein aggregates throughout the hippocampus and subiculum (54.6 ± 20.2 mm^2^).

We first used a ‘top-down’ approach to determine alignment with previous observations of hippocampal changes in AD (Mrdjen et al., 2019). We found that the hippocampus from the CN participant held Aβ plaques and PHF1-TAU NFTs-NTs, albeit less than CIND and ADD hippocampus. Interestingly, proximity analysis showed that microglia interacted heavily with pathologic Tau, particularly in CA1 region. These microglia harbor increasing signatures of possible ‘disease-associated microglia’ (DAM) in ADD (Guo et al., 2020; Keren-Shaul et al., 2017). We found similar associations using bottom-up data-driven approach for microglial and pathologic Tau. Surprisingly, microglia interactions with Aβ immunoreactive structures were less frequent, potentially indicating microglia respond more to pathologic Tau. Our assessment of using the data-driven, bottom-up approach uncovered unique proteopathy and cellular spatial associations. The Aβ AP-region appeared more dominant in the CIND sample relative to the ADD sample, which is dominated by the pathologic Tau TT-region, perhaps a reflection of the established correlation of tauopathy with dementia in AD. The composition of N- and G-regions are particularly interesting as they represent regions of tissue that are mostly free from proteopathy and are found even in ADD. The exact relationship between proteopathy and glial lineages in the continuum of AD is unclear and could be regulated by several elements. For instance, immunogenicity towards Aβ or Tau oligomers in CN and CIND could vary from ADD samples (Brunello et al., 2020; Li and Selkoe, 2020; Thomas et al., 2012), or ADD may have a more reactive microglial-dependent immune response relative to those present in the CN and CIND hippocampus (Sáez et al., 2006).

The neuronal features that were imaged here are from those surviving neurons that have endured AD, including non-proteopathy bearing (PHF1-TAU and Aβ negative) and proteopathy-bearing neurons (PHF1-TAU or Aβ positive). We identified several molecular signatures that could have contributed to neuron survival. First, as in many animal models and recently in human tissue, we found pathologic Tau to be localized (but not exclusive) to neuronal synapses (Lemke et al., 2020; Phongpreecha et al., 2021). *Drosophila* to mouse studies have shown that synaptic PHF1-TAU interferes with synaptic vesicles (Sahara et al., 2014; Tai et al., 2014; Zhou et al., 2017). Second, we observed Aβ accumulation proximal to synaptic signal of glutaminergic and GABAnergic neurons where others have suggested acute Aβ exposure can lead to GABAnergic imbalance and altered neurotransmission (Li and Selkoe, 2020; Quevenco et al., 2019; Wang et al., 2017).

Interestingly, we also observe potential compensatory mechanisms, including ubiquitin proteosome system (UPS) and mitochondrial bioenergetics (i.e., MFN2, Figure 6) across all individuals, but most prominently in the person with ADD. Here, we detected K48-linked ubiquitin in proteopathy rich neurons, in particular preferential enrichment in NFTs-NTs. Similar enrichment of K48-linked ubiquitin has been reported in older AD brains (Nakayama et al., 2019). The role of preferential K48-ubiquitination on PHF1-TAU oligomers is still unclear; however, others have proposed a “soak-up” mechanism to sequester the toxic effect of protein aggregates and promote UPS mediated clearance (Arrasate et al., 2004; Mund et al., 2018). Importantly, we also found a prevalence of MFN2 expression in proteopathy-burdened hippocampal regions, particularly in areas of pathologic Tau. Here MFN2, an outer membrane mitochondria fusion protein, could maximize the oxidative capacity of the disease-free neurons, a potential response to a stressor associated with accumulation of pathologic Tau. MFN2 also inhibits mitochondrial fission, preventing autophagy, and thus helping to maintain mitochondria for the surviving neurons (Han et al., 2020; Sita et al., 2020; Wang et al., 2009). Given the role of neuron-mitochondrial dysfunction found in multiple models of AD, MFN2 may represent an important target for maintaining survival in proteopathy-susceptible neuron populations. Taken together, our bottom-up data-driven approach of studying disease neighborhoods mapped out phenotypic changes as a function of proteopathy and identified potential coping mechanisms of hippocampal pyramidal neurons to withstand the stressors of proteopathy-associated AD pathogenesis. Although we focus on neuronal cell neighborhoods here, the unsupervised approach can be extended to map the response of proteostatic stress on glial and vascular cells. Targeted approaches for the other key homeostatic populations (microglia, astrocytes, and oligodendrocytes) are imperative in understanding the effect of cellular crosstalk (feedback and feedforward), heterogeneity in activity, population abundance, and localization to disease and cognitive status. The framework we have created here could be applied to investigate CNS cellular biology in all these scenarios.

Like many neurodegenerative diseases, AD is phasic, undergoing biochemical changes that induce cellular stress and remodeling of anatomical neighborhoods. Detailed mapping of the biochemical, cellular, and architectural changes can help develop strategies for dynamic models of disease, potentially from a limited number of patient samples. Future studies on larger patient cohorts could order individuals, based on the spatial features we have derived here, along disease pseudo-time as means to quantify disease progression as reflected in spatio-molecular changes. Pseudo-time ordering has been used for fate mapping, tumor progression, hematopoietic development, and to map the layered and cell-specific degeneration of hippocampal sclerosis (Bendall et al., 2014; Cid et al., 2021; Kimmey et al., 2019; Lang et al., 2019; Loeffler-Wirth et al., 2018; Weinreb et al., 2020). Inferred disease staging can help sort the temporal function of biochemical and molecular regulators in AD clinical progress, identify founding factors in AD, and uncover disguising factors that potentially play roles for neurological resilience in ‘healthy’ individuals. Such modelling of multiplex imaged human brain tissue may help understand whether Aβ42 oligomers followed by PHF1-TAU oligomers are primers or followers to degeneration, or if other underlying bad molecular actors precedes proteopathy (e.g., aberrant immune activation).

MIBI-TOF and our study also had limitations. First, because MIBI-TOF imaging scans physically sputter material off the imaged sample, it is thus destructive. Still, thin sections can be re-scanned multiple times and serial sections can be used for further experimentation. Second, this is a probe-based technology that largely relies on antibody availability and specificity for detection of limited target antigens (∼40), though the staining and integration of serial sections with different antibody panels can exponentially increase the N-dimensionality of probing factors. Still, probe-based technologies also tend to be more robust and quantitative compared to *de novo* techniques that offer a wider, albeit variable, array of multiplexing. Although unexplored here, as an elemental mass spectrometry method, MIBI-TOF can simultaneously quantify presence of essential and non-essential metals, such as Fe, Cu, Mn, Zn, and Al, along with the N-number proteins which can contribute to AD pathogenesis (Chen et al., 2016). Third, findings are contingent upon the rigor of patient and postmortem tissue characterization (i.e., clinical diagnosis and neuropathological assessment), and tissue integrity. Dynamic measurement of postmortem human brain tissue is not trivial, requiring a large number of carefully annotated samples from people of varying ages. Nevertheless, inferences of AD progression can be made with limited postmortem sample via use of systematic characterized subjects, tissue, multiplex approaches, and data-driven modelling. Compatibility, as we have shown here, with the archival (i.e., FFPE) format these tissues are typically stored in is critical for successful interrogation.

Our work paves the way for the development of novel approaches for quantitative, simultaneous, multiplexed imaging in human neuropathology. Utilizing the wealth of RNA sequencing data, potential disease mechanisms driving cell-cell and cell-proteopathy interactions can be isolated and now imaged at the proteomic level in their native tissue with our multiple, high-resolution MIBI-TOF platform. Combining multiple spatial biomolecular technologies (e.g., MALDI, RNA-ISH, super-resolution microscopy) will enable even deeper understanding of biomolecular interplay in neurodegeneration. Integration of this spatially multiplexed data with further clinical histories will drive insights on possible cellular mechanisms underlying symptomatic disease progression or resilience in CNS disease.

## ACKNOWLEDGMENTS

We would like to thank Dr. Rosario Sanchez-Pernaute for her critical inputs on the manuscript. Humphrey Vijayaragavan-Bossé and Teddy Cannon-Merchant for their unconditional love and moral support. Dr. Edward Fox for his assistance with the brain tissue repository. This work was supported by NIH grants R01 AG056287, R01 AG057915, R01 AG068279, U19 AG065156, UH3 CA246633 and P30 AG066515. N.F.G. was supported by NCI CA246880-01 and the Stanford Graduate Fellowship. Canadian Institute of Health Research Postdoctoral Fellowship (J.P.O.) T32 AI007290 to B.J.C and E.F.M. D.M. was supported by Schweizerischer Nationalfonds zur Förderung der Wissenschaftlichen Forschung (P2ZHP3_181563), Novartis Foundation for medical-biological Research (#18B067), and the Glenn Foundation for Medical Research Postdoctoral Fellowship in Aging Research.

## STAR METHODS

### KEY RESOURCE TABLE

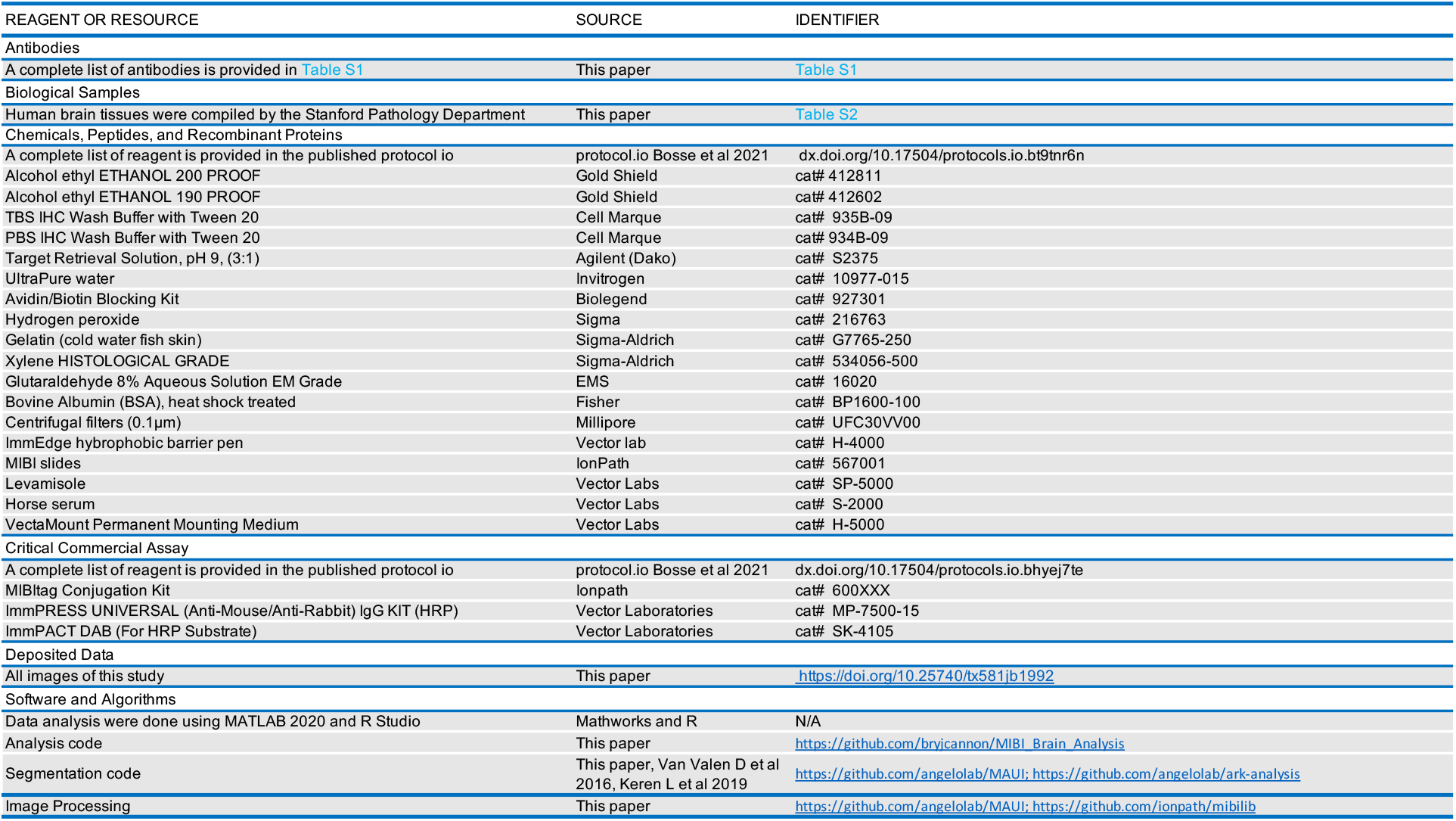

### CONTACT FOR REAGENTS AND RESOURCE SHARING

Further information and request for reagents and resource should be directed to and will be fulfilled by the Lead Contact, Sean Curtis Bendall.

### EXPERIMENTAL MODEL AND SUBJECT DETAILS

#### Human sample acquisition and patient consent

Human brain samples were obtained from Stanford Pathology Department. Average post-mortem interval of tissues ranges between 2.9-15 hours. Clinical diagnosis and neuropathological scores were generated by Stanford clinicians and pathologist. The clinical diagnosis was based on DSM-IV criteria. AD neuropathologic change and severity scores were evaluated by NIA-AA guidelines (Hyman et al., 2012; Montine et al., 2011). All three cases (CN, CIND and ADD) also were evaluated for neuropathologic evidence of vascular brain injury, Lewy body disease, or hippocampal sclerosis by NIA-AA guidelines and those with any evidence of these three neuropathologic changes were excluded. Neuropsychological test battery results for CN participants within 1 year of death were in the upper three quartiles for the research cohort.

### MIBI-TOF EXPERIMENTAL DETAILS

#### Antibody conjugation

Primary antibodies, reporter isotopes, and titers can be found in Table S1. Metal conjugation of primary antibodies were prepared as described previously (Bendall et al., 2013) and is detail in dx.doi.org/10.17504/protocols.io.bhyej7te (Bosse et al., 2021a). Post conjugation antibodies were diluted in Candor PBS Antibody Stabilization solution (Candor Bioscience GmbH, Wangen, Germany) to 0.1 mg/ml, and stored at 4°C for short-term use and for long-term used were lyophilized using the following protocol dx.doi.org/10.17504/protocols.io.bhmgj43w (Camacho et al., 2021).

#### MIBI staining

FFPE tissue blocks of whole hippocampus or brain TMA were sectioned using a microtome. The 5μm thick tissue sections were mounted onto silanized-gold slides for MIBI-TOF staining. Gold slides containing tissue were baked at 70°C overnight. Tissue sections were then processed and stained with a cocktail of antibodies is as detailed in dx.doi.org/10.17504/protocols.io.byzrpx56 (Bosse et al., 2021b). Stained tissue slides were preserved under vacuum until required for MIBI-TOF scan.

#### IHC staining

For IHC validation of primary antibody target specificity, FFPE human hippocampus or brain TMA, of 5μm thickness, were sectioned onto standard glass slide. Slides containing tissue were baked at 70°C overnight. Tissue sections were then processed and stained using the Sequenza method with single post-metal conjugated primary antibody. Staining method is detailed in dx.doi.org/10.17504/protocols.io.bf6ajrae (Bosse et al., 2021c)

#### MIBI-TOF image acquisition

Spectral images of stained hippocampus or brain TMA tissue were obtained using the MIBI-TOF mass spectrometer equipped with a Hyperion ion source. Xenon primary ions were used to sputter the stained tissue sample at doses ranging from average 17.9nAmp ms pixel^2^/μm^2^ for 500μm^2^ to 52.3nAmp ms pixel^2^/μm^2^ for 400μm^2^ FOVs. MIBI-TOF parameters used in acquiring all imaging data is detailed in Table S3.

#### TOF calibration and spectral image generation

Post-acquisition, time of flight was calibrated using sodium (Na+ 22.99 amu) and gold (Au+ 196.97amu). Counts for each mass were determined by the start and end of their respective spectral peaks. For instance, in CD56 antibody conjugate to Nd145, the counts for masses between 144.7 to145.2 were used and so on. Spectral images for each FOV were generated by converting the mass-spec pixel data to multidimensional tiffs.

#### Image processing (background subtraction and denoising)

After extracting raw data, images were background removed, noise corrected, and tiled to form a composite image of the entire issue section. Background (Bg) signal and noise were removed using a streamline in house developed graphic user interface(gui) in Matlab (https://github.com/angelolab/MAUI). Bg signal highly correlate with Au channel from the slide, on the edges and holes in the tissue. The Bg signal (T=0.1-0.2, gaussian kernel with R =1 pixel) were removed from all channels. High frequency signal was considered noise from MIBI-TOF brain spectral data. Also, many signals from brain tissue, especially those from synaptic markers (SYN, PSD95, VGLUT1/2, VGAT), CD56, and CD47 are dense signals. These dense signals were used as the high frequency cut-offs for our Fast Fourier transform (FFT) approach to filter noise from all brain spectral images. Furthermore, signal spill over due to adducts and oxides, were compensated for the following conjugates: pASyn to Abeta, IBA1 to PSD95, GFAP to CD105, MBP to MCT1, MCT1 to CD33, TAU to MFN2, TH, MAP2, pTDP43, 8OHG and Aβ. Processed data were stored as image Tiff files and .mat files for data processing and analysis.

#### Image stitching and normalization for visualization

Image stitching for visualization purposes was performed using custom jython scripts within the Fiji/ImageJ image processing environment and utilizing existing plugins. Individual tiles first underwent a flat-field correction (FCC) within tissue masked areas. Specifically, tissue masks were generated using the thresholded C12 channel, as well as the inverse of thresholded Au197 and Empty139 channels. The FCC reduces the effects of mass dependent aberrations within the fields of view. The FCC is performed by summing, down sampling, and smoothing each channel from each field of view. This smoothly varying intensity map was then used to normalize each individual field of view channel. Additionally, to correct for spatially uneven detection, each tile was cropped by 1% on each side to eliminate edge effects, followed by an additional FCC. This second FCC generates an intensity map for each tile by averaging, down sampling, and smoothing all channels, the result of which should average out biologically distinct distributions, leaving only an intensity response map. These maps are then respectively used to normalize each field of view’s channels. The “Grid stitching” plugin then stitched these tiles together, with overlaps specified by the acquisition parameters. Each channel was then smoothed using a 2-pixel median filter, and linearly auto scaled to maximize the visual contrast for each channel. For visualization with MIBItracker, each image was down sampled to have the largest dimension equal to 2048 pixels. For many channels it was possible to reduce the gridded pattern artifact via bandpass filtering in Fourier space. This is acceptable as biological information is typically not aligned in this way. However, channels whose signal varied significantly over a low frequency across the imaged area were not bandpass filtered, as these components would also be removed. Related to this work, tiling functionalities are now available through the Ionpath’s github repository https://github.com/ionpath/mibilib.

Additional stitching was done via a custom Matlab tiling script linking start and end coordinates of each scanned images.

### DATA PROCESSING AND DISCRETIZATION ANALYSIS

#### Global expression pattern

Background removed and denoised image TIFs (one file per marker) containing the pixel intensities were used to determine the global expression levels per TIF file. Each pixel represents marker expression intensity in a 500 or 400 μm^2^ area and where the variances of marker co-expression between different areas of the human brain.

#### Cellular segmentation

Nuclei segmentation was carried out using an adapted version of DeepCell (Keren et al., 2018; Moen et al., 2019; Valen et al., 2016). *Training data*. Epithelial cell nuclei stained with dsDNA or HH3 were used as core training data as described previously (Keren et al., 2018). *Segmentation of brain images*. All images (841 TIFs of HH3) were subjected to the trained network and were normalized in the same fashion as the training images, by subtracting an averaging filter and dividing by the std of the whole image. *Post processing*. Probability map for the ‘nuclear interior’ were thresholded and separated to generate nuclear masks as described before (Keren et al., 2018). Cell border expansion on these nuclear masks were performed to partially capture cell bodies. Nuclear morphology information including, area, circularity, eccentricity, major and minor axis lengths (*regionprops*, MATLAB 2019b) were used to infer cell type between neuronal and glial cells due to their large differences in cell versus nucleus sizes. To calculate mean marker expression values for each segmented cell, the TIFs containing the pixel expression values were combined with the expanded nuclear segmentation masks. A data frame containing information of each segmented cell, with an allocated cellular ID (row), is populated with the corresponding values of each marker expression (column). In addition, supporting overlay files that track the spatial information for each cell in each FOV were created. To linearize the data and deal with extreme outliers, all mass channels were log-transformed in addition to 99^th^ quantile normalization in in RStudio (Version 2.1)

#### Object segmentation

Thus far, we have used nuclear segmentation and blanket marker expression data in our analysis. To enable the use of object structures, such as plaques and tangles, we had to employ EZSegmenter – a MATLAB *regionprops* thresholding-based segmentation GUI developed in-house and available as a part of the MIBI image processing toolkit here: https://github.com/angelolab/MAUI. Multiplexed TIF images from multiple FOVs are loaded into the GUI. Channel select displays signal intensities of MIBI images and are used to determine object masks. Additional settings (e.g. Gaussian Blur, minimum and maximum object pixel size) are fixed across FOVs. Masks are then used to extract pixel – level signal intensities across each channel and then cell size normalized before import into an output cell table csv file. In addition to segmenting plaques, tangles, and vessels, we were able to improve cell segmentation by including parts of cells that had their nuclei mechanically destroyed by the tissue preparation process or were not scanned by the MIBI. All parameters for each objects segmented can be found in Table S3B. This new data underwent similar normalization steps as the original segmented cell data.

#### Manual Gating ADD hippocampus (1024 × 1024 images)

Akin to classical single cell analysis in cytometry, stepwise manual gating strategy starting from the lowest to highest mean pixel levels was carried out. We first gated on CD31+, CD105+ and MCT1+ to label the endothelial fraction, the remaining fraction were gated for IBA1+ and CD45+ to classify microglia, and subsequently GFAP+ was used to label the astrocytes. The final remaining meta fraction was labeled as neurons and oligodendrocytes. Gated populations were overlayed back onto the spectral images and incorrect labels were curated using a multi-channel imaging platform (Mantis viewer, Parker Institute, https://mantis.parkerici.org/).

#### Single object clustering & phenotyping

##### 512 × 512 CN, CID, & ADD Images

Initial gating was performed using common marker panels. Endothelial CD31, MCT1, CD105: endothelial cells; CD45 and IBA1: microglia; GFAP: astrocytes

##### 1024 × 1024 ADD Images

To assign a lineage for each segmented cell, a sequential gating strategy was performed on the transformed marker expression, starting with CD31, MCT1, CD105 for endothelial cells; then CD45 and IBA1 for microglia; then GFAP for astrocytes. Remaining ungated cells were then assigned as neurons. The accuracy of manual gating was curated by a multichannel imaging application MantisViewer. The curated expression data was imported back into R for UMAP analysis, heatmaps and object co-proximity analysis. Masks of neurons identified by cellular segmentation were merged with masks of the object segmented data (microglia, astrocytes, endothelia, plaques, and tangle-threads) to obtain a comprehensive repertoire of neuronal and non-neuronal cell types, and proteopathy features.

#### Object co-proximity analysis

Thresholds for the co-proximity analysis were empirically determined by calculating the shortest physical distance of the periphery of a cell to another cell or proteopathy feature in a pairwise manner, and in a predefined pixel-radius. A distance interaction matrix was calculated for each pixel-radius, for 25(≈10μm), 50(≈20μm), 100(≈39μm), 150(≈59μm), 200(≈78μm), 250(≈98μm) and 300(≈117μm) pixel-radius, and the optimal distance used for thresholding was based on counts and frequencies of object pairwise interaction. The threshold of t = 25-pixel-radius for NVU, t = 50-pixel radius for tangle-threads and t = 100-pixel radius for amyloid plaques. The null hypothesis (Ho): all objects do not interact with each other and are equidistantly dispersed throughout each FOV. The baseline object interaction (Ho) frequencies were calculated via the number of objects that would appear within threshold radius (t) by chance. The random chance was estimated by taking a proportion of objects appearing within the area of the thresholded radius and the entire FOV. Since this analysis was performed on individual FOVs, the ratio of the observed neighbor counts versus their estimated counterparts were used to normalize the values between FOVs.

#### Software, data & code availability

Software for running the MIBI-TOF equipment was developed by SAI (MiniSIMS 2 Data Systems Version 5.5.4.0). All the data described in this work, including channel images, segmentation masks and cellular and object identities can be accessed through a web interface and downloaded at https://mibi-share.ionpath.com. The code for the analysis can be downloaded at https://github.com/bryjcannon/MIBI_Brain_Analysis. All the information required for cell segmentation including the manual training data, trained neural networks, and code for training and running DeepCell (now incorporated into a cell segmentation known as MESMER) are available at Ark-Analysis https://github.com/angelolab/ark-analysis.

