## Supplemental Info for "Single-cell Spatial Proteomic Imaging for Human Neuropathology": Supplemental_Figures.pdf

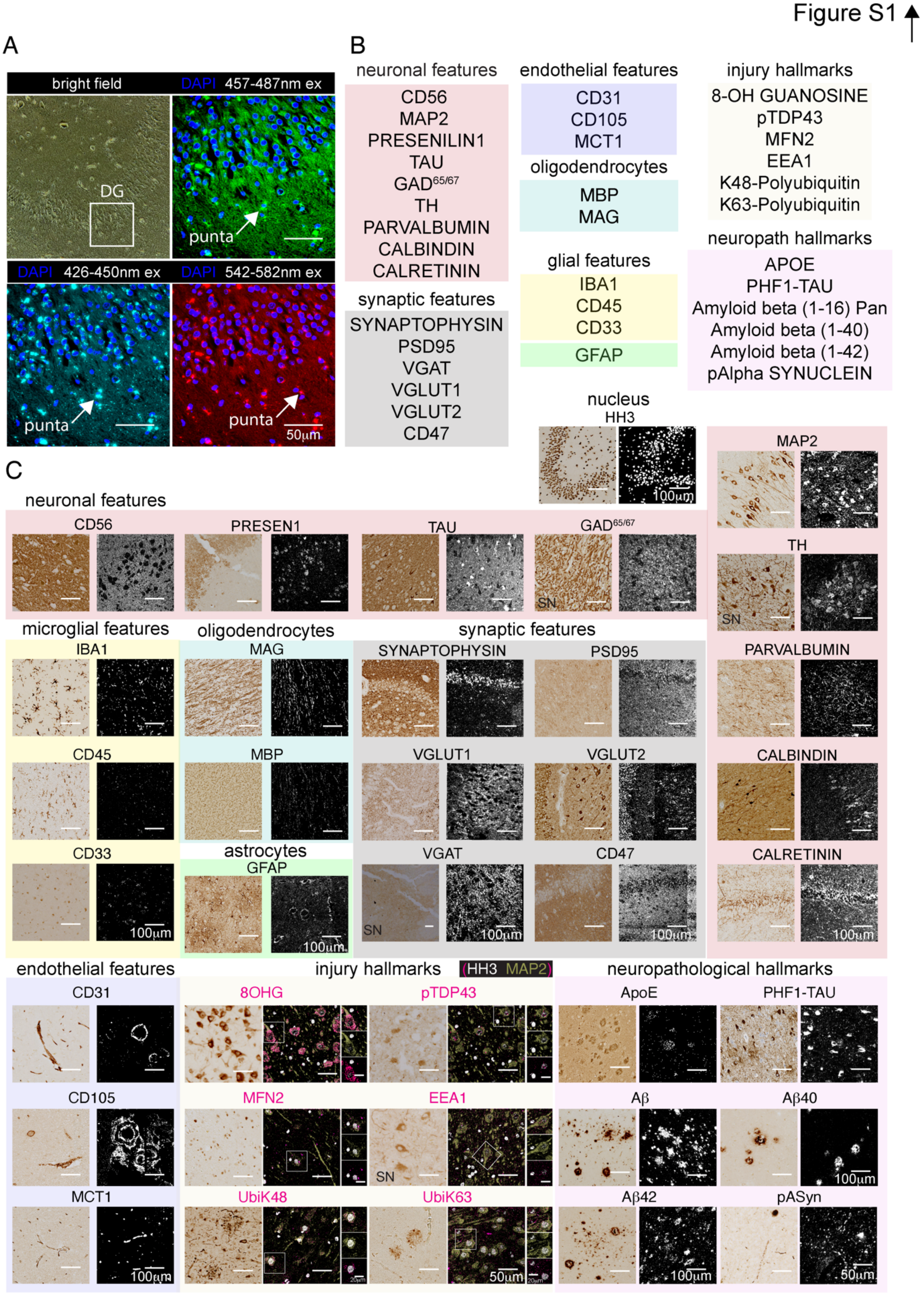
Figure S1
A
bright field
DAPI 457-487nm ex
DG
punta
DAPI 426-450nm ex
DAPI 542-582nm ex
50µm
B
neuronal features
CD56
MAP2
PRESENILIN1
TAU
GAD65/67
TH
PARVALBUMIN
CALBINDIN
CALRETININ
synaptic features
SYNAPTOPHYSIN
PSD95
VGAT
VGLUT1
VGLUT2
CD47
endothelial features
CD31
CD105
MCT1
oligodendrocytes
MBP
MAG
glial features
IBA1
CD45
CD33
GFAP
injury hallmarks
8-OH GUANOSINE
pTDP43
MFN2
EEA1
K48-Polyubiquitin
K63-Polyubiquitin
neuropath hallmarks
APOE
PHF1-TAU
Amyloid beta (1-16) Pan
Amyloid beta (1-40)
Amyloid beta (1-42)
pAlpha SYNUCLEIN
nucleus
HH3
100µm
C
neuronal features
CD56
PRESEN1
TAU
GAD65/67
SN
MAP2
TH
PARVALBUMIN
CALBINDIN
CALRETININ
microglial features
IBA1
CD45
CD33
oligodendrocytes
MAG
MBP
astrocytes
GFAP
synaptic features
SYNAPTOPHYSIN
PSD95
VGLUT1
VGLUT2
VGAT
CD47
100µm
endothelial features
CD31
CD105
MCT1
injury hallmarks
(HH3 MAP2)
8OHG
pTDP43
MFN2
EEA1
UbiK48
UbiK63
50µm
20µm
neuropathological hallmarks
ApoE
PHF1-TAU
Aβ
Aβ40
Aβ42
pASyn

**Figure S1. Validation of MIBI-TOF antibody panel on FFPE archival brain tissue.**
(A) Autofluorescence photomicrographs of an unstained human hippocampus section under bright field, green (457-487nm excitation), cyan (426-450nm excitation) and red (542-582nm excitation) fluorescence channels. (B) List of brain specific targets used for MIBI-TOF analysis. (C) IHC and MIBI-TOF antibody validation and cross comparison of chromogenic and spectral images.

Figure S2

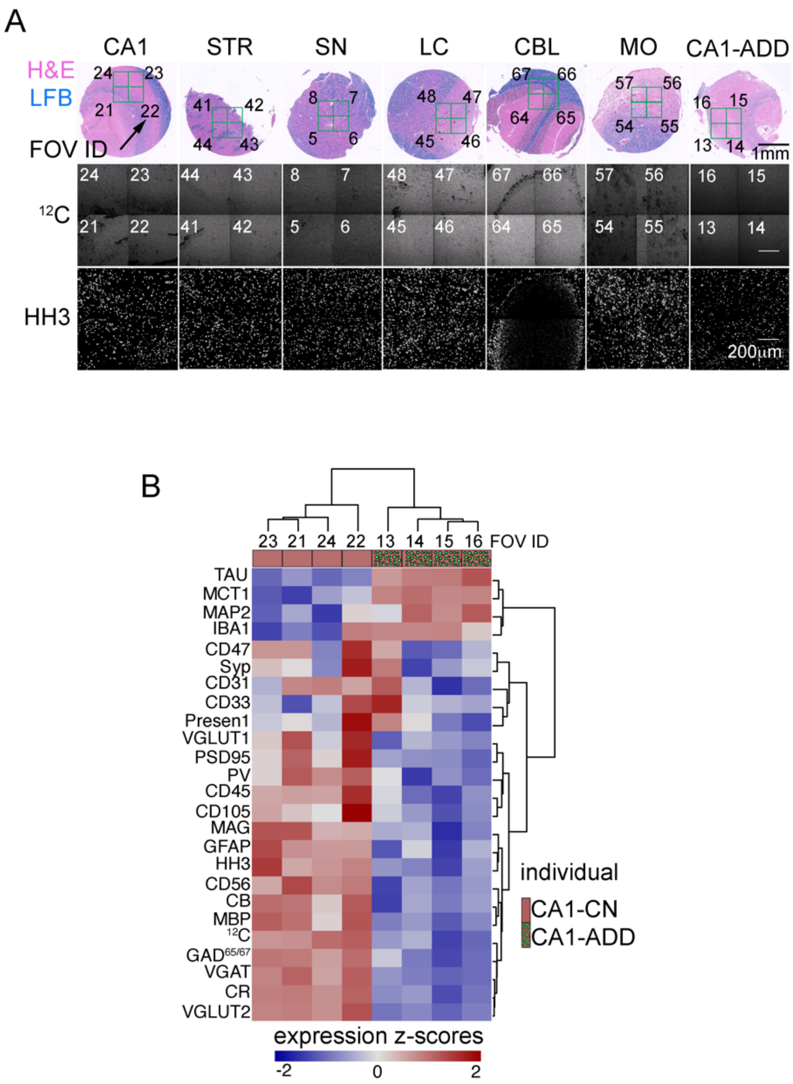
A
CA1
STR
SN
LC
CBL
MO
CA1-ADD
H&E
LFB
FOV ID
1mm
12C
HH3
200µm
B
FOV ID
TAU
MCT1
MAP2
IBA1
CD47
Syp
CD31
CD33
Presen1
VGLUT1
PSD95
PV
CD45
CD105
MAG
GFAP
HH3
CD56
CB
MBP
12C
GAD65/67
VGAT
CR
VGLUT2
individual
CA1-CN
CA1-ADD
expression z-scores
-2
0
2

**Figure S2. Global Expression Analysis of Multiplex Proteomic Data Stratifies cognitive normal and ADD CA1 cores.**

(**A**) Photomicrograph of brain TMA cores stained with LFB/H&E to reveal myelin and other cellular structures (top). Green boxes depict the rastered area (4x4, 500μm$^2$) and each FOV is assigned an ID. The $^{12}C$ spectral images in grey scale unveil field uniformity between each FOV of the same brain region (middle), and spectral images of HH3 show the presence and distribution of nuclei (bottom). (**B**) Heatmaps showing the average pixel z-score distribution of proteins per FOV. Variance between and among FOVs are stratified in an unsupervised manner into normative and disease. Columns and rows are hierarchically clustered (Euclidean distance).

Figure S3

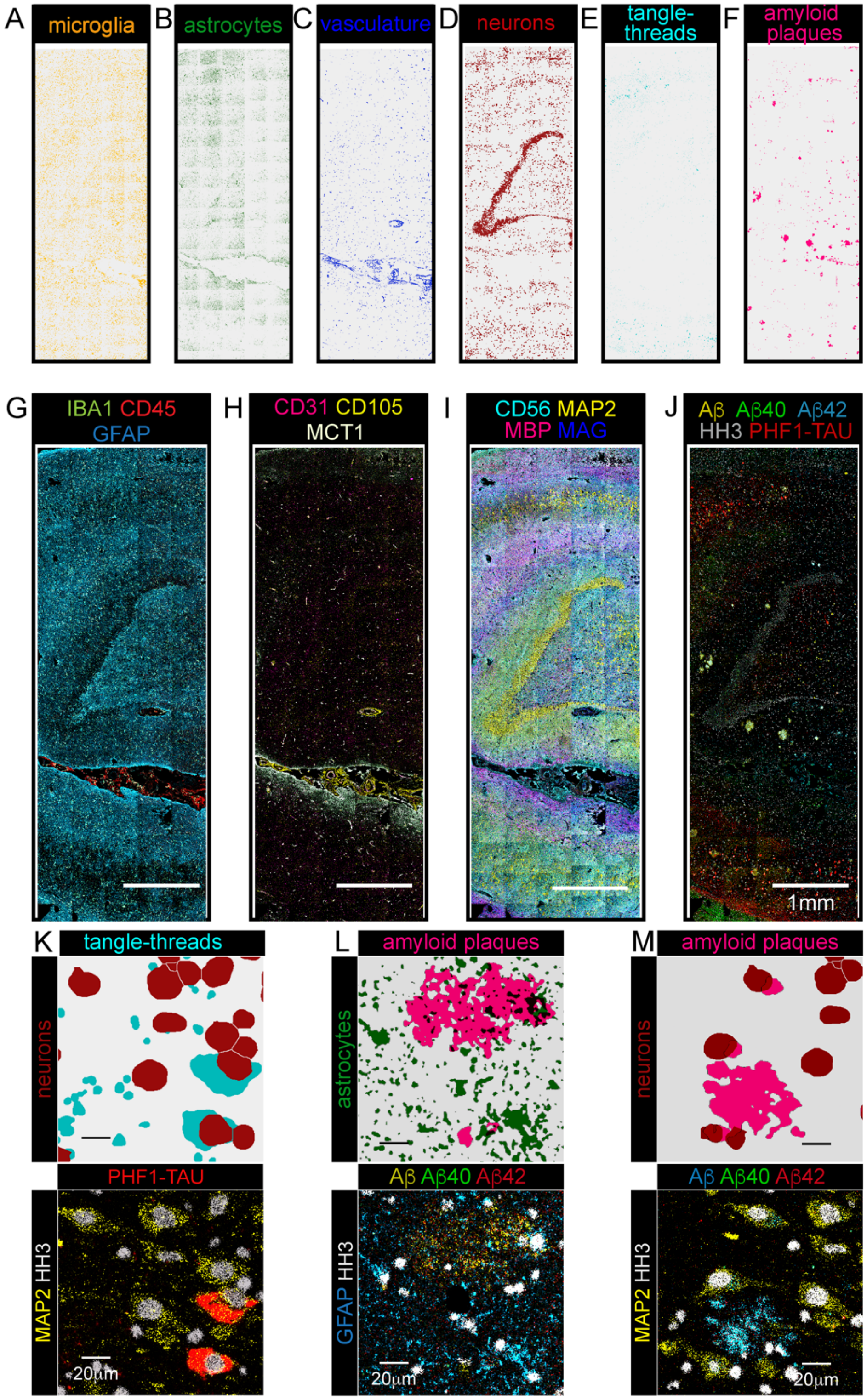
A
microglia
B
astrocytes
C
vasculature
D
neurons
E
tangle-threads
F
amyloid plaques
G
IBA1 CD45 GFAP
H
CD31 CD105 MCT1
I
CD56 MAP2 MBP MAG
J
Aβ Aβ40 Aβ42 HH3 PHF1-TAU
1mm
K
tangle-threads
neurons
PHF1-TAU
MAP2 HH3
20μm
L
amyloid plaques
astrocytes
Aβ Aβ40 Aβ42
GFAP HH3
20μm
M
amyloid plaques
neurons
Aβ Aβ40 Aβ42
MAP2 HH3
20μm

**Figure S3. Cell Phenotype Maps (CPM) of Segmented features in ADD hippocampus.**
**(A-F)** Individual Cell Phenotype Maps (CPM) of nuclear and object segmented masks labelled by their phenotypes. **(G-J)** MIBI-TOF pseudo-colored spectral images for the different cells which are identified by their pan-markers (GFAP: astrocytes, IBA1-CD45: microglia, CD31-CD105-MCT1: vessels, MAP2: neurons, CD45: immune cells). **(K-M)** Segmented mask overlays of cell and proteopathy phenotypes with corresponding MIBI-TOF pseudo-colored spectral images.

Figure S4A-C ↑

cognitively normal no dementia (CN)

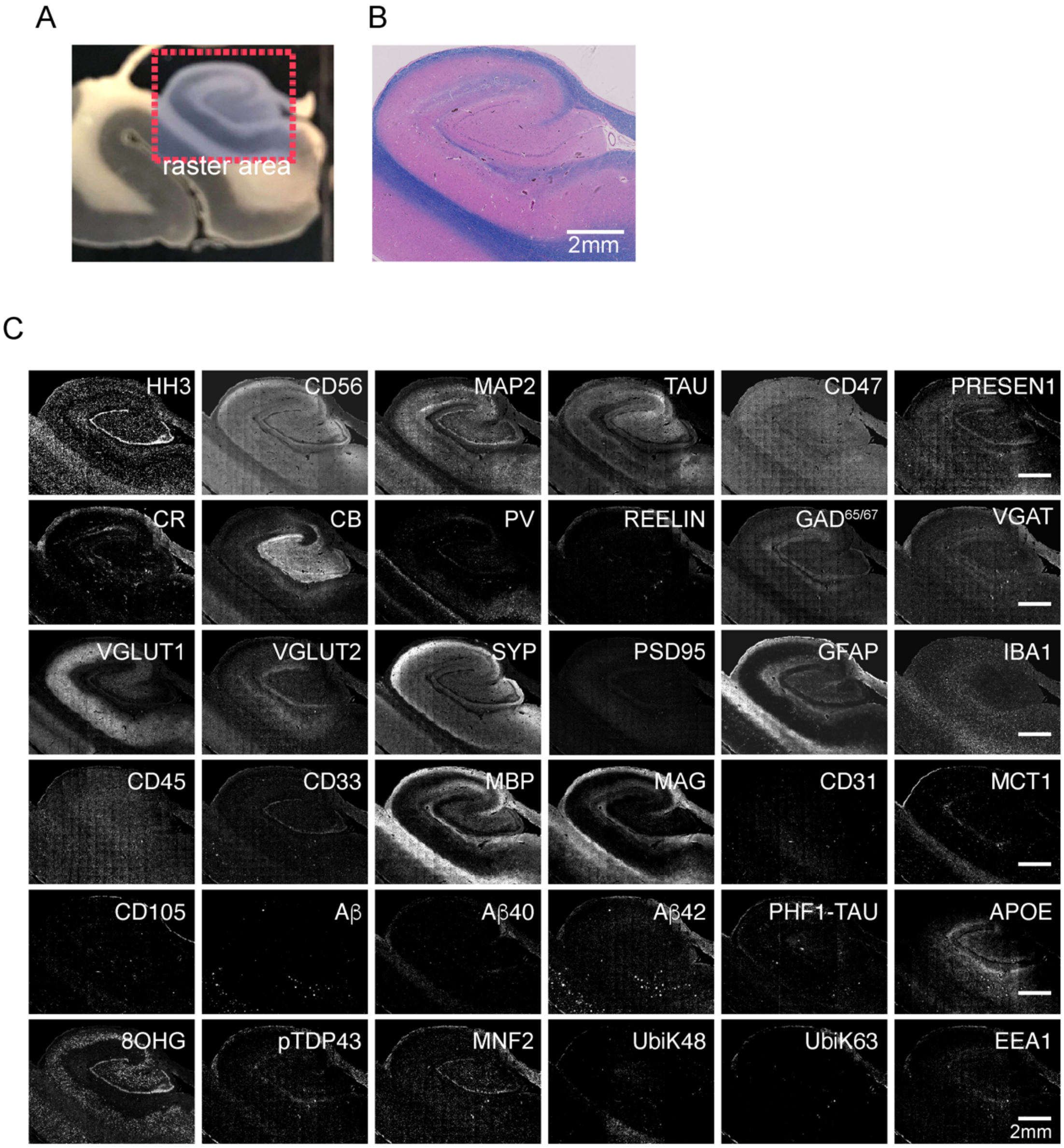

Figure S4D-F

cognitively impaired no dementia (CIND)

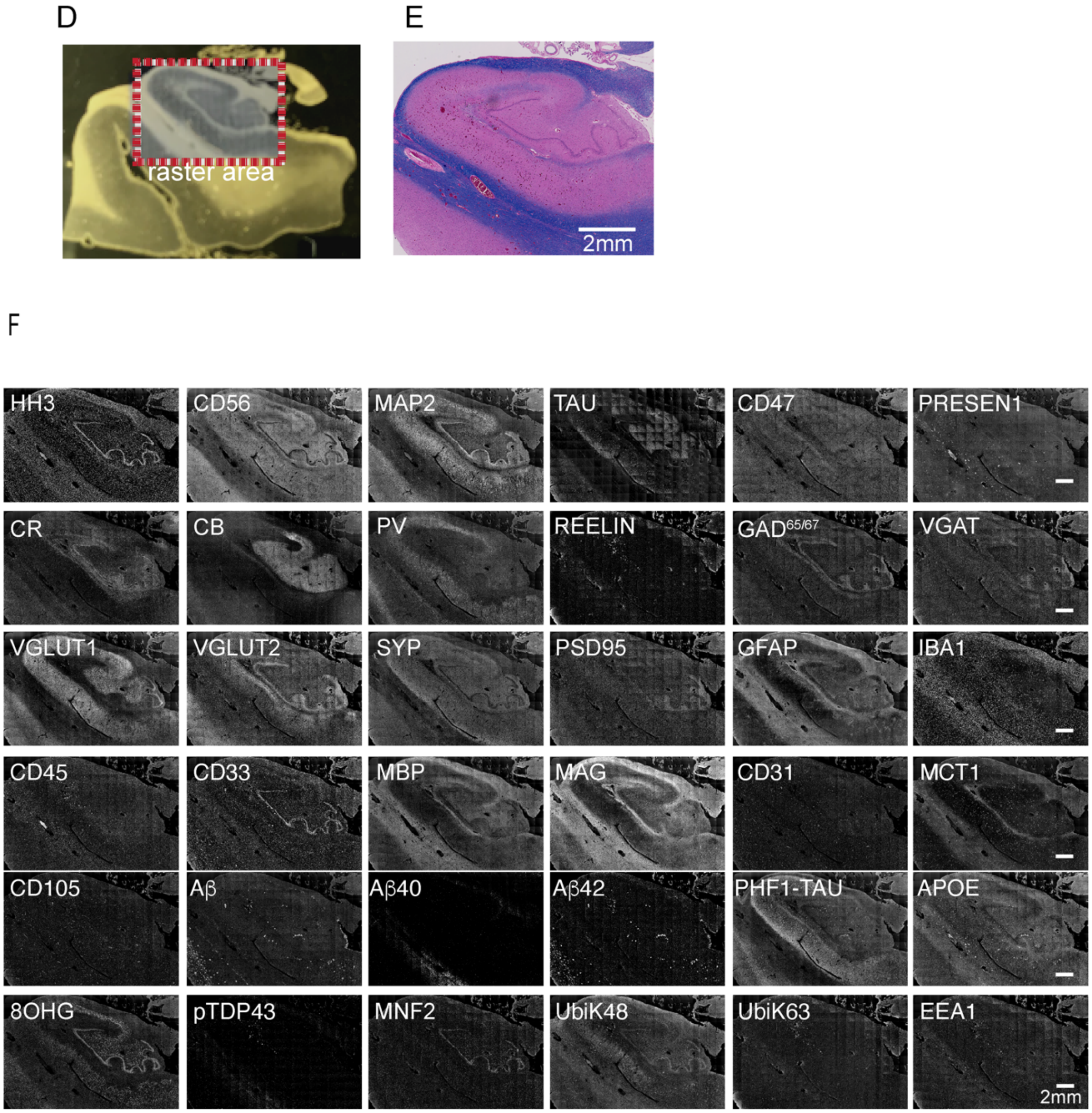

Figure S4G-I

cognitively impaired with dementia (ADD)

G

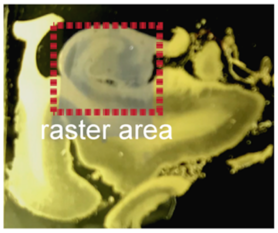


H

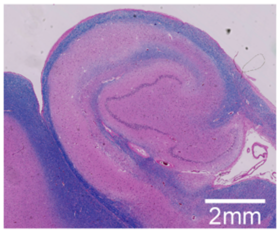


I

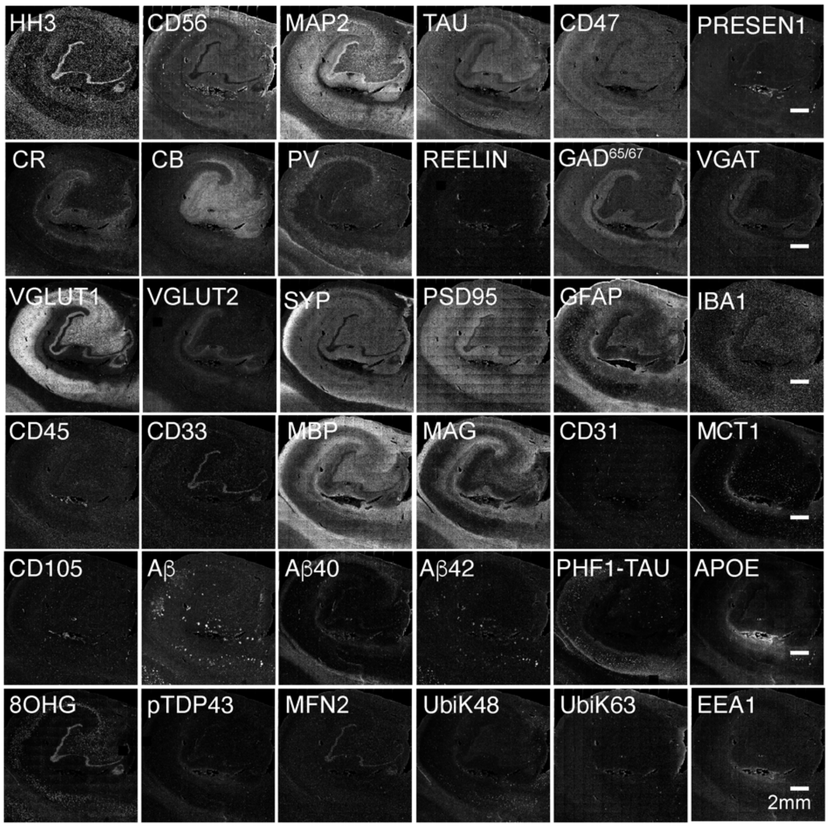

Figure S4J-L

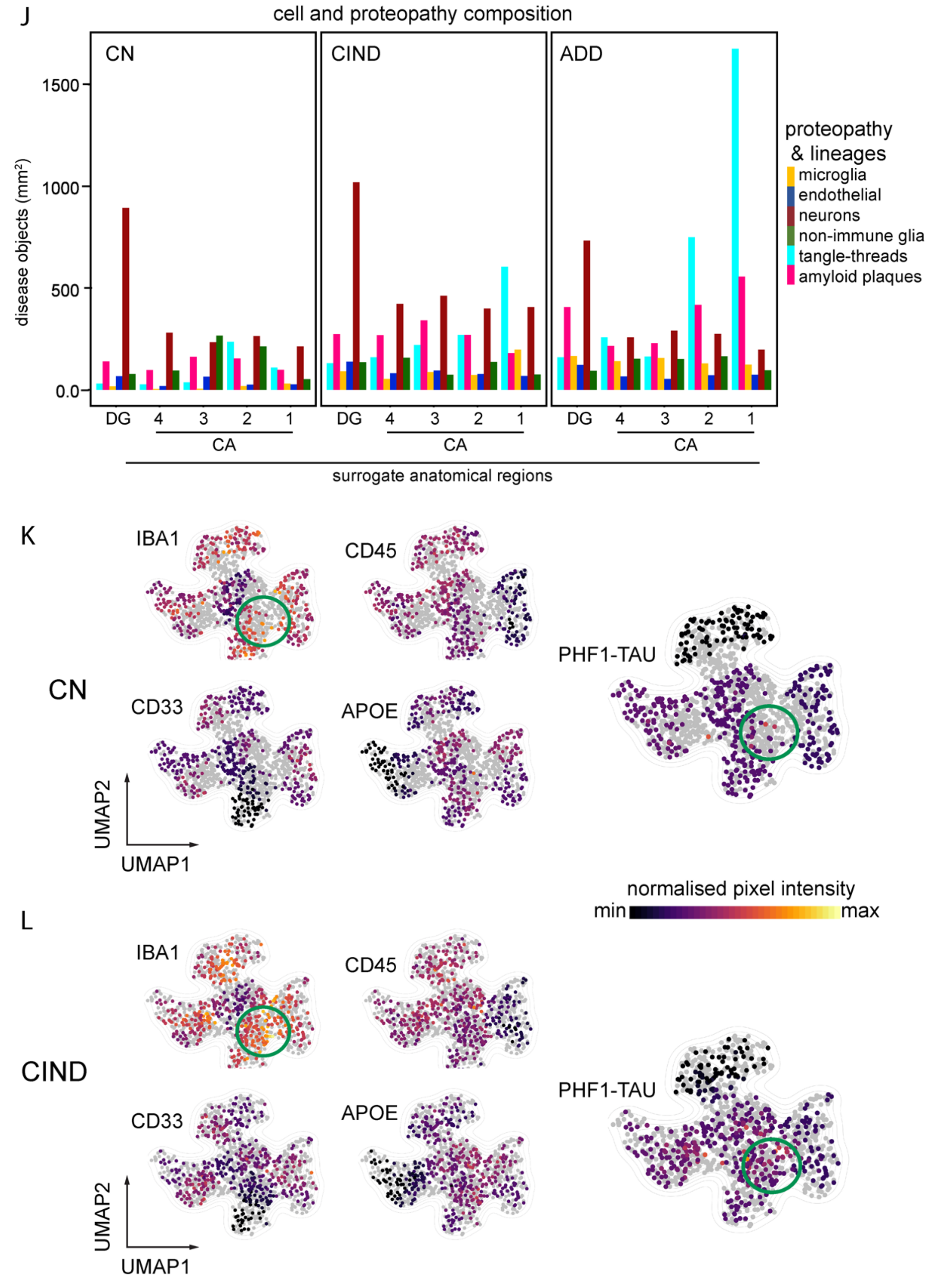


**Figure S4. Single-plex spectral images of CN, CIND and ADD and their breakdown of cell and proteopathy characteristics in anatomical regions.**

**(A, D, G)** Archival mid-section hippocampi from CN, CIND, or ADD on conductive gold-slides. Red dashed boxes show rastered regions respectively. **(B, E, H)** Serial sections stained with standard histological Luxol Fast Blue (LFB) for myelin with hematoxylin and eosin (H&E) for cells. **(C, F, I)** Single plex MIBI spectral images for all 36 targets that were simultaneously acquired. Images for CN is of 300 FOVs tiled; CIND is of 260 FOVs tiled, ADD is of 196 tiled FOVs. Each FOV is 500μm$^2$. Images are gray scaled. **(J)** Normalized counts of cells and proteopathies within each mm$^2$ of tissue across each hippocampal anatomical region and individual. **(K)** UMAP projections of CA1 microglia phenotyping channels in CN. Gray represents CA1 microglia from ADD, CIND samples. **(L)** UMAP projections of CA1 microglia phenotyping channels in CIND. Gray represents CA1 microglia from ADD, CN samples. Projection calculated using phenotypic markers. CIND and ADD are subsampled so that all conditions are represented by 447 microglia, the total number of CA1 microglia in the CN condition.

Figure S5

A

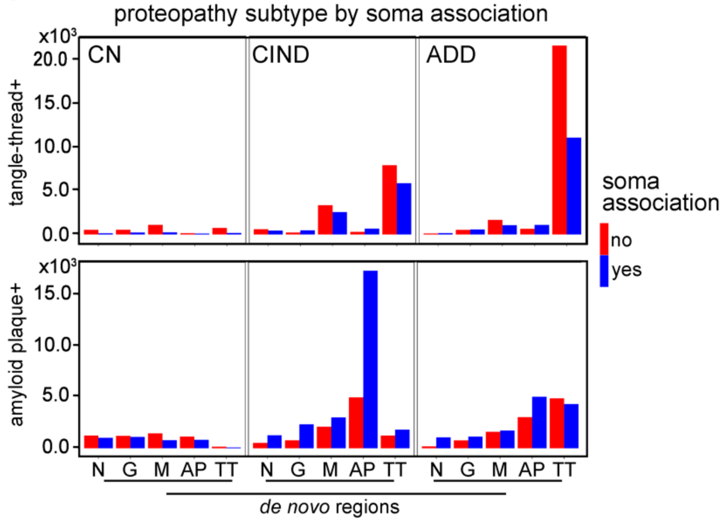
proteopathy subtype by soma association
x10³
CN
CIND
ADD
tangle-thread+
amyloid plaque+
20.0
15.0
10.0
5.0
0.0
soma association
no
yes
N G M AP TT
de novo regions

**Figure S5. Breakdown of cell and proteopathy characteristics in de novo regions.**
**(A)** Raw counts of proteopathies associated with a cell soma or not across the de novo regions. Highest counts of plaques found in AP region within CIND (majority soma associated), highest counts of tangles found within TT in ADD (majority not soma associated). **(B)** Normalized mean expression of non-immune glia, microglia, endothelial cells, and neurons with associated phenotyping channels. Broken down across the *de novo* regions.

Figure S6

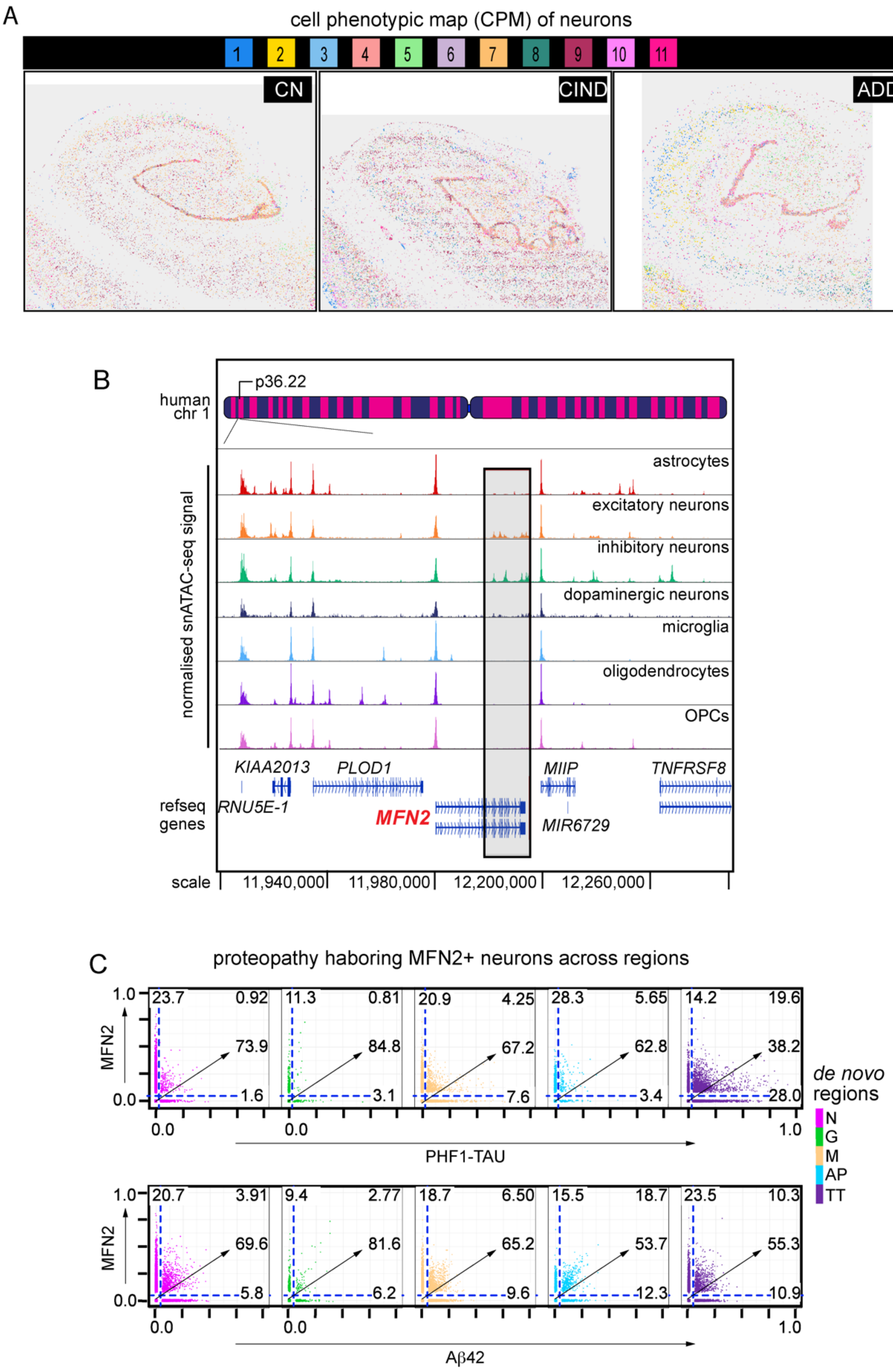
A
cell phenotypic map (CPM) of neurons
1
2
3
4
5
6
7
8
9
10
11
CN
CIND
ADD
B
p36.22
human chr 1
normalised snATAC-seq signal
astrocytes
excitatory neurons
inhibitory neurons
dopaminergic neurons
microglia
oligodendrocytes
OPCs
KIAA2013
PLOD1
MIIP
TNFRSF8
RNU5E-1
MFN2
MIR6729
refseq genes
scale
11,940,000
11,980,000
12,200,000
12,260,000
C
proteopathy haboring MFN2+ neurons across regions
MFN2
1.0
0.0
23.7
0.92
73.9
1.6
11.3
0.81
84.8
3.1
20.9
4.25
67.2
7.6
28.3
5.65
62.8
3.4
14.2
19.6
38.2
28.0
PHF1-TAU
de novo regions
N
G
M
AP
TT
20.7
3.91
69.6
5.8
9.4
2.77
81.6
6.2
18.7
6.50
65.2
9.6
15.5
18.7
53.7
12.3
23.5
10.3
55.3
10.9
Aβ42

**Figure S6. Neuronal sub-clustering and MFN2 specificity to neurons.**
**(A)** Overlay of tissue samples with neurons alone, colored by subcluster assignment. **(B)** ATAC-seq data from human brain tissue showing localized MFN2 open peaks in neuron cell types only. Gray box represents open MFN2 peaks in neurons alone. **(C)** MFN2 and proteopathy channel relationships across all de novo regions.
